# Statistical testing and power analysis for brain-wide association study

**DOI:** 10.1101/089870

**Authors:** Weikang Gong, Lin Wan, Wenlian Lu, Liang Ma, Fan Cheng, Wei Cheng, Stefan Grünewald, Jianfeng Feng

## Abstract

The identification of connexel-wise associations, which involves examining functional connectivities between pairwise voxels across the whole brain, is both statistically and computationally challenging. Although such a connexel-wise methodology has recently been adopted by brain-wide association studies (BWAS) to identify connectivity changes in several mental disorders, such as schizophrenia, autism and depression [Cheng et al., 2015a,b, 2016], the multiple correction and power analysis methods designed specifically for connexel-wise analysis are still lacking. Therefore, we herein report the development of a rigorous statistical framework for connexel-wise significance testing based on the Gaussian random field theory. It includes controlling the family-wise error rate (FWER) of multiple hypothesis testings using topological inference methods, and calculating power and sample size for a connexel-wise study. Our theoretical framework can control the false-positive rate accurately, as validated empirically using two resting-state fMRI datasets. Compared with Bonferroni correction and false discovery rate (FDR), it can reduce false-positive rate and increase statistical power by appropriately utilizing the spatial information of fMRI data. Importantly, our method considerably reduces the computational complexity of a permutation-or simulation-based approach, thus, it can efficiently tackle large datasets with ultra-high resolution images. The utility of our method is shown in a case-control study. Our approach can identify altered functional connectivities in a major depression disorder dataset, whereas existing methods failed. A software package is available at https://github.com/weikanggong/BWAS.

## 1 Introduction

The human brain connectome is usually modelled as a network. In the brain’s network, accurately locating the connectivity variations associated with phenotypes, such as clinical symptoms, is critical for neuroscientists. With the development of neuroimaging technology and an increasing number of publicly available datasets, such as the 1000 Functional Connectomes Project (FCP) [Biswal et al., 2010], Human Connectome Project (HCP) [Glasser et al., 2016] and UK Biobank [Miller et al., 2016], large-scale, image-based association studies have become possible and should help us improve our understanding of human brain functions.

Using a priori knowledge of brain parcellation (e.g. AAL [Rolls et al., 2015]) or an adoption of data-driven parcellation (e.g. ICA [Beckmann and Smith, 2004]) to analyze the human connectome is the most popular approach, and many statistical methods have been designed for them [Zalesky et al., 2012; Kim et al., 2014]. However, with the availability of large datasets, increasing the spatial specificity in the functional connectivity analysis should provide a deeper insight into the brain connectome. Therefore, in this paper, a statistical framework for brain-wide association study (BWAS) is proposed [Cheng et al., 2015a,b, 2016]. It directly uses *voxels* as nodes to define brain networks, and then tests the associations of each *functional connectivity* with phenotypes.

To conduct a systematic, fully-powered BWAS, two main issues should be carefully addressed. First, a multiple correction method to control the false-positive rate of massive univariate statistical tests should be developed. Second, a power analysis method to estimate the required sample size should be designed. One may ask whether the methods widely used in region-level studies can be directly generalized to connexel-level studies. Two issues hinder such direct generalization. First, the statistical tests have more complex spatial structures in BWAS. Therefore, as shown in our analysis, some widely-used multiple correction methods which do not utilize the spatial information of data (e.g. Bonferroni correction and false discovery rate (FDR) [Benjamini and Hochberg, 1995; Benjamini and Yekutieli, 2001]) may not be powerful enough to detect signals. Second, although non-parametric permutation methods [Nichols and Holmes, 2002] may account for the complex structures among hypothesis tests to provide a valid threshold, they are computationally very expensive in connexel-wise studies, owing to the requirement of performing billions of statistical tests. Therefore, an accurate and efficient method for multiple comparison problem and power analysis is needed.

Random field theory (RFT) is an important statistical tool in brain image analysis, and it has been widely used in the analysis of task fMRI data and structure data [Ashburner and Friston, 2000]. Statistical parametric maps (SPM) are usually modelled as a discrete sampling of smooth Gaussian or related random fields [Penny et al., 2011]. The random field theory can control the FWER of multiple hypothesis testings by evaluating whether the observed test statistic, or the spatial extent of clusters exceeding a cluster-defining threshold (CDT), is large by chance, which is known as peak-level and cluster-level inference respectively. Since Adler’s early work on the geometry of random field [Adler, 1981; Adler and Taylor, 2009], theoretical results for different types of random fields have been obtained, such as the Gaussian random field [Friston et al., 1994; Worsley et al., 1996b], the *t*, *χ*^2^, *F* random fields [Worsley, 1994; Cao, 1999], the multivariate random field [Taylor and Worsley, 2008], the cross-correlation random field [Cao et al., 1999]. Among them, only the cross-correlation field is designed for connectivity analysis. In that framework, the voxel-level functional connectivity network is modelled as a six-dimensional cross-correlation random field, and the maximum distribution of the random field is used to identify strong between-voxel connections. Different from the above works, the aim of BWAS is to identify connectivities that are associated with phenotypes. To our knowledge, no previous works have addressed this problem. In this paper, we show that the statistical map of BWAS, under the null hypothesis, can be modelled as a Gaussian random field with a suitable smoothness adjustment. Therefore, topological inference methods, such as peak intensity and cluster extent, are generalized from voxel-wise analysis to functional connectivity analysis.

Besides controlling the type I error rate, estimating power or the required sample size for BWAS is also important. In genetics, for example, a high-quality GWAS analyzing one million single nucleotide polymorphism (SNP) usually requires tens of thousands of samples to reach adequate statistical power. In contrast, previous BWAS analyses of schizophrenia, autism and depression have only had sample sizes less than one thousand [Cheng et al., 2015a,b, 2016]. Therefore, compared to GWAS, it is natural to ask if BWAS, which is usually based on a limited sample size, can withstand the rigors of a large number of hypothesis tests. In this regard, most existing power analysis methods are designed for voxel-wise fMRI studies, including, for example, the simulation based method [Desmond and Glover, 2002], the non-central distribution based method [Mumford and Nichols, 2008], and the method based on non-central random field theory (ncRFT) [Hayasaka et al., 2007]. Among them, the ncRFT-based method can both take into account the spatial structure of fMRI data and avoid time consuming simulation. Therefore, to analyze the power of BWAS, we adopted a methodology similar to that of the ncRFT-based method [Hayasaka et al., 2007]. The signals at functional connectivities are modelled as a non-central Gaussian random field, and the power is estimated by a modified Gaussian random field theory.

In this paper, a powerful method to address the multiple comparison problem is proposed for BWAS (Figure 1). This method uses Gaussian random field theory to model the spatial structure of voxel-level connectome. It can test the odds that either the effect size of every single functional connectivity (peak-level inference) or the spatial extent of functional connectivity clusters exceeding a cluster-defining threshold (cluster-level inference) is large by chance. The performance of the method is tested in two resting-state fMRI datasets, and in both volume-based and surface-based fMRI data. Our method can control the false-positive rate accurately. Compared with Bonferroni correction and false discovery rate (FDR) approaches, our method can achieve a higher power and filter out false-positive connectivities by utilizing the spatial information. In addition, we develop a modified Gaussian random field theory to explicitly approximate the power of peak-level inference (Figure 2). Power can be estimated in any specific location of connectome efficiently, which can help to determine the sample size for BWAS. The utility of our method is shown by identifying altered functional connectivities and estimating the required sample sizes in major depression disorder. The software package for BWAS can be downloaded at https://github.com/weikanggong/BWAS.

**Figure 1:**
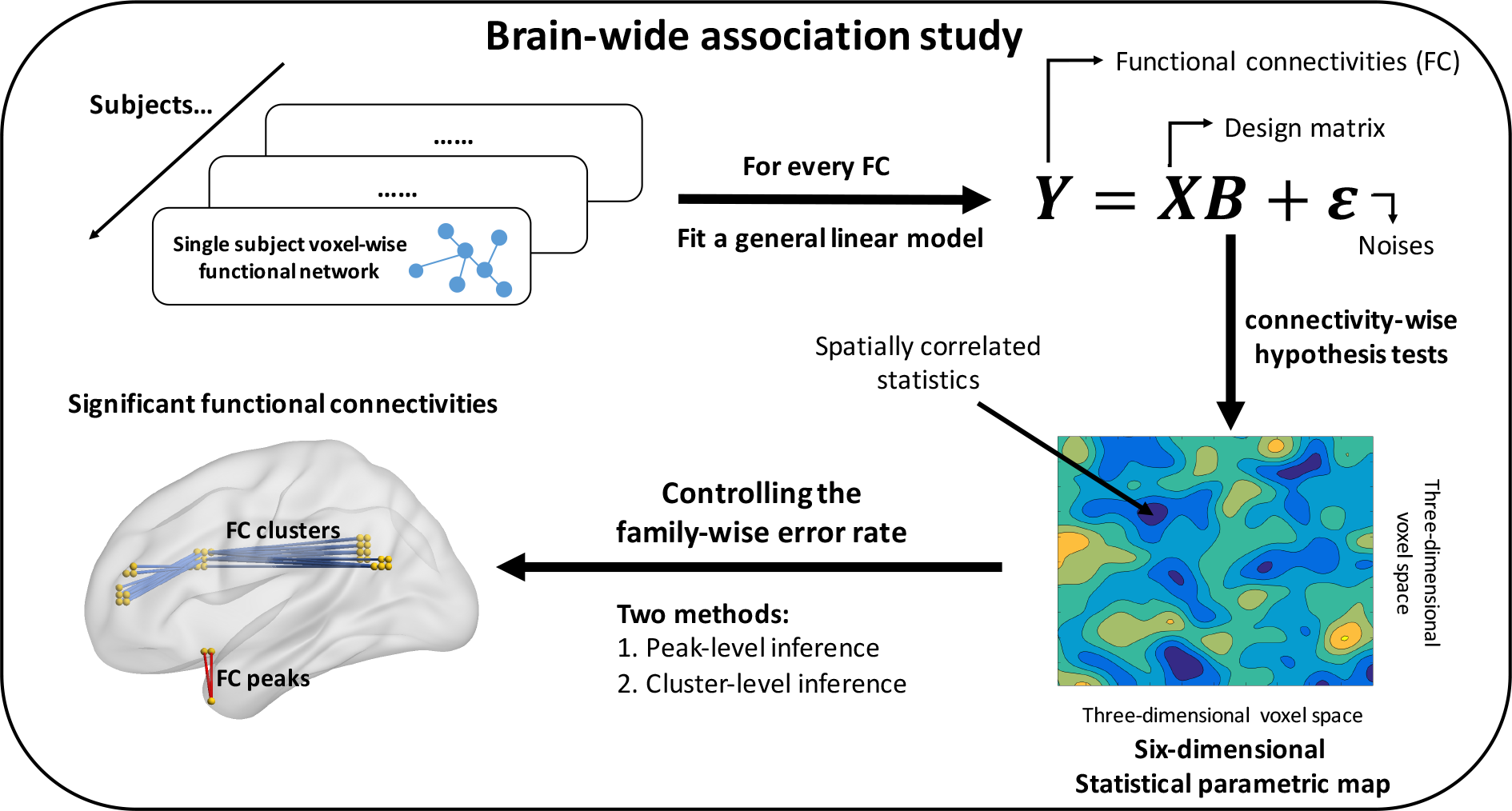
A flow chart of brain-wide association study. First, we estimate the voxel-level brain network for each individual. Then, we perform connectivity-wise statistical tests to test the association between each functional connectivity and a phenotype of interest. Finally, peak- and cluster-inference approaches are used to identify significant signals.

**Figure 2:**
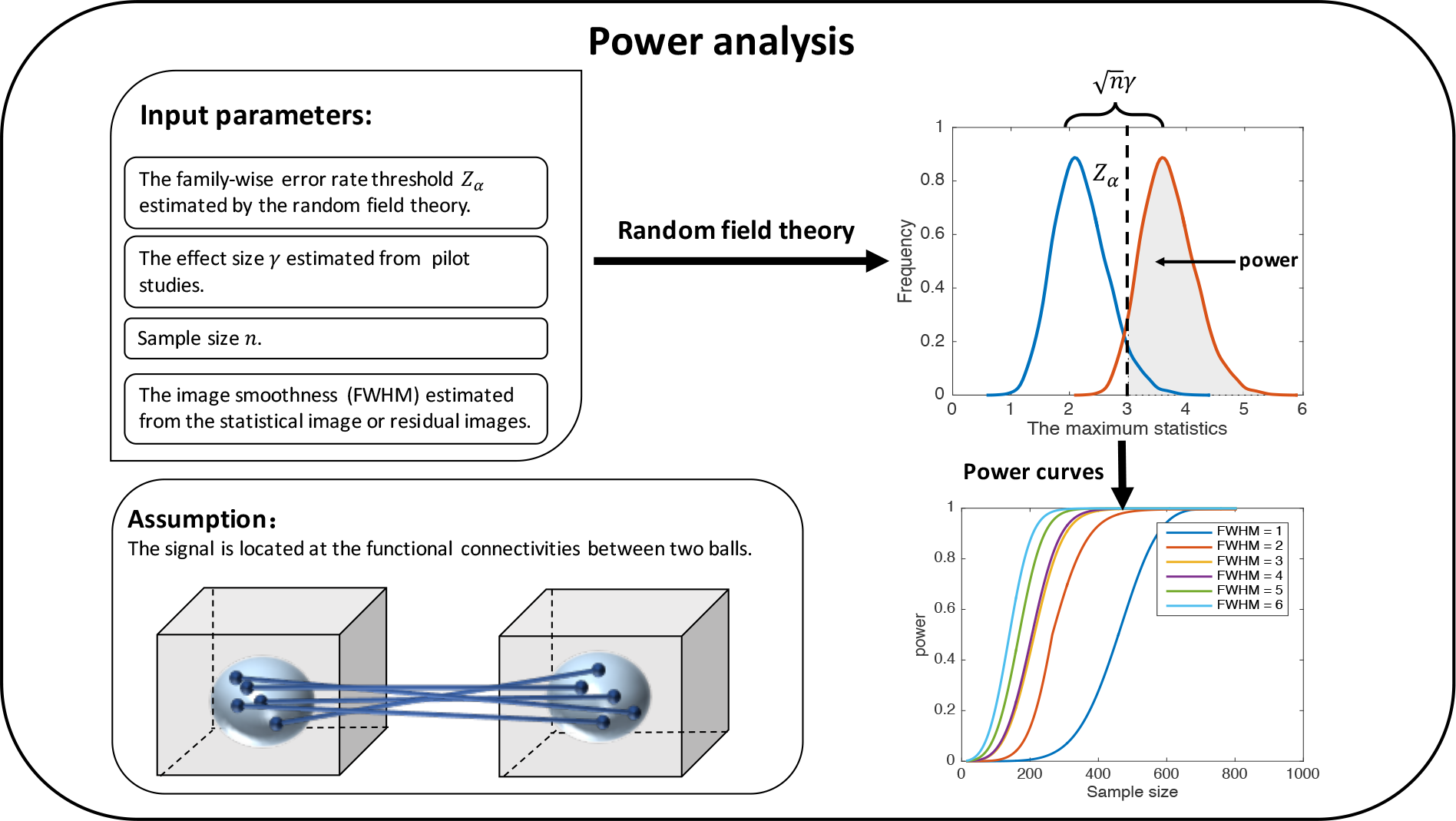
Power analysis for brain-wide association study. To estimate power, we first calculate the FWER-corrected threshold *Z_α_* of peak-level inference, and then estimate the effect size *γ* from a prior statistical map of BWAS. For a target sample size *n*, and a smoothness level FWHM, we can estimate the power using the random field theory, which is defined as the probability of finding at least one true-positive signal in a region, in which the false-positive rate *α* is controlled at a certain level in the whole search region. Finally, the power under different sample sizes and smoothness levels can be estimated iteratively.

## 2 Materials and Methods

### 2.1 Connexel-wise general linear model

The popular general linear model approach is used in BWAS. Briefly, a voxel-level functional network is estimated for each subject using the fMRI data, and the association between each functional connectivity and phenotype of interest is tested using the general linear model.

In detail, the individual functional network is constructed first by calculating the Pearson correlation coefficients (PCC) between every pair of voxel time series. Let *m* be the number of voxels, *s* be the subject, and 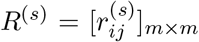 be the *m* × *m* functional network matrix for subject *s*. Each element of *R*^(*s*)^ is the correlation coefficient between voxel time series *i* and *j* for subject *s*. An element-wise Fisher’s Z transformation is then applied as 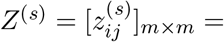 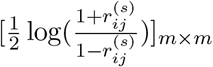, so that 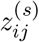 will approximate a normal distribution. For every functional connectivity, a general linear model (GLM) is fitted by

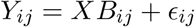

where, 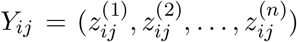 is an *n* × 1 vector of functional connectivities between voxel *i* and *j* across *n* subjects, *X* is the common *n* × *q* design matrix, 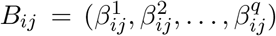 is a *q* × 1 vector of regression coefficients, and *∊_ij_* is an *n* × 1 vector of random error, which is assumed to be an independent and identically distributed Gaussian random variable 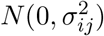 across subjects. The ordinary least square estimator for *B_ij_* is 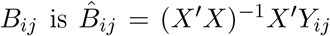, and for 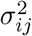, it is 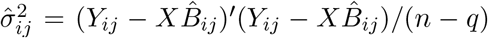. Then, a Student’s t-statistic at functional connectivity between voxel *i* and *j* can be expressed as:

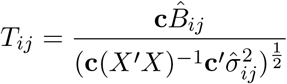

where **c** is a 1 × *q* contrast vector. In BWAS, let 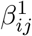 be the primary variable of interest, and 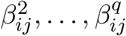 be the nuisance covariates included in the regression model. The contrast **c** = (1, 0,…,0) will be used to test the hypothesis 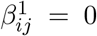, and the *T_ij_*-statistics will reflect the significance of the primary variable. Other contrasts can also be used depending on the study design. Finally, the Student’s t random variable at each functional connectivity *T_ij_* is transformed to a Gaussian random variable *Z_ij_*by transforming *T*-statistics to *p*-values and then to *Z*-statistics.

After the above steps, the connexel-wise Z-statistics form a statistical parametric map in a six-dimensional Euclidian space. The reason is that the spatial location of each Z statistic (or functional connectivity) can be uniquely represented by the coordinates of its two endpoints, each of which is a voxel in a three-dimensional space. Therefore, the structure of the statistical map can be modelled by the random field theory, and the topological inference methods for multiple hypothesis testings are developed in the subsequent Section.

### 2.2 Multiple comparison correction using topological inference methods

#### 2.2.1 Peak-level inference

Peak-level inference controls FWER among multiple hypothesis testings on functional connectivities, i.e., the probability of finding at least one false-positive functional connectivity is controlled under certain level α. It is assumed, under the null hypothesis, that the statistical parametric map of BWAS is a discrete sampling of smooth and stationary Gaussian random fields with mean zero and variance one. To control the FWER of multiple hypothesis testings, the maximum distribution of the random field should be known. In our paper, its tail distribution is approximated by the expected Euler characteristic (EC) of the excursion set of random field. The detailed derivation is given in the Appendix. We sketch an overview of the result here.

Let *Z*(*p*, *q*), *p* ∈ 𝒫, *q* ∈ 𝒬 be a (P+Q)-dimensional Gaussian random field spanned by a P-dimensional random field 𝒫 and a Q-dimensional random field 𝒬. At high threshold *z*_0_, its maximum distribution has a general form [Adler and Taylor, 2009; Worsley et al., 1996b]:

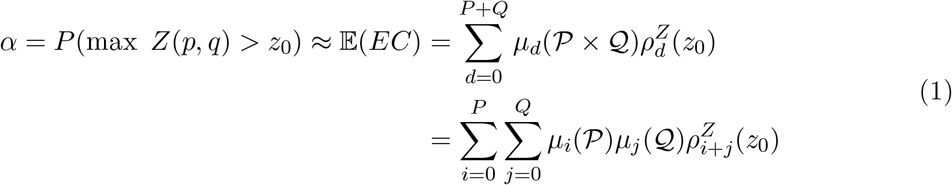

where the *μ_d_*(·) is the *d*-th dimensional intrinsic volume of the random field, and 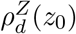 is the *d*-th dimensional EC-density for the Gaussian random field at threshold *z*_0_ (*z*_0_ > 0). The method for calculating *μ_d_*(·) and 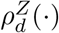 are shown in the Appendix. Therefore, the *α*-level FWER-corrected threshold *z*_0_ can be found using equation (1), and for one-tailed tests, functional connectivities with Z-values larger than *z*_0_ (or smaller than −*z*_0_) are declared as significant.

For different kinds of BWAS analysis, P and Q in (1) can take different values. For example, for the widely-used volume-based fMRI data, we use P=Q=3 (Result Section 3.2.1). If the connectivities are estimated between pairwise vertices on cortical surface, we use P=Q=2 (Result Section 3.2.1), and if the connectivities are estimated between subcortical structures and cortical surface, we use P=3 and Q=2. The estimated FWER-corrected threshold is usually less conservative than Bonferroni correction method, because the intrinsic volume *μ*_d_(·) in equation (1) takes into account both the number of hypothesis tests performed and the correlations among tests, and an increasing of spatial smoothness can make the FWER-corrected threshold *z*_0_ lower. For BWAS, the equation (1) can be approximately estimated using the results of Gaussian random field, provided that the spatial smoothness is estimated correctly. The reason is that the statistical map of BWAS is generated by a series of non-linear transformation of original fMRI images. As a result, we calculate equation (1) as:

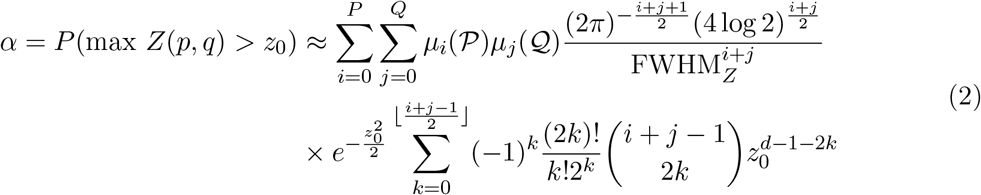

where FWHM*_Z_* is the adjusted full-width at half maximum of the Gaussian smooth kernel, which is a function of the original smoothness of fMRI images. A proof of the equation (2) and smoothness estimation approach are shown in the Appendix.

#### 2.2.2 Cluster-level inference

Cluster-level inference is also popular in brain image analysis. Here, inference is based on the observed cluster size exceeding certain cluster-defining threshold (CDT) [Friston et al., 1994]. We are usually interested in whether the observed cluster size is large by chance, i.e., where the size is on the upper tail of the distribution of maximum cluster size under the null hypothesis. We show that, similar to the voxel clusters in three-dimensional space, the functional connectivity cluster (FC cluster) can also be defined rigorously. Its size can be used as a test statistic for statistical inference, and it has a clear interpretation.

A voxel cluster is a set of spatially connected voxels. To define a FC cluster, we first illustrate the neighbourhood relationship between two functional connectivities. Let the endpoints of two functional connectivity be (*x*_1_, *y*_2_) and (*x*_2_, *y*_2_), if their endpoints are non-overlapped voxels, then two functional connectivities are neighbours if both *x*_1_, *x*_2_ and *y*_1_, *y*_2_ are spatially adjacent voxels. If they share a same endpoint (e.g. *x*_1_ = *x*_2_), then they are neighbours if *y*_1_, *y*_2_ are spatially adjacent voxels. Some examples of FC neighbours are shown in Figure 3A. Now, consider an undirected graph 𝒢 with *k* nodes, where the nodes are k functional connectivities and two nodes are connected if they are neighbours, then these *k* functional connectivities form a FC cluster if they form a connected component in the graph 𝒢. Some examples of FC clusters are shown in Figure 3B. There are five voxel clusters A, B, C, D, E in a two-dimensional image. The FCs between AB, BC and AD are different FC clusters, and FCs within voxel cluster E also form a single FC cluster. An algorithm for finding FC clusters can be implemented based on the above definition. In our analysis, We use Dulmage-Mendelsohn decomposition to find connected components in graph 𝒢.

**Figure 3:**
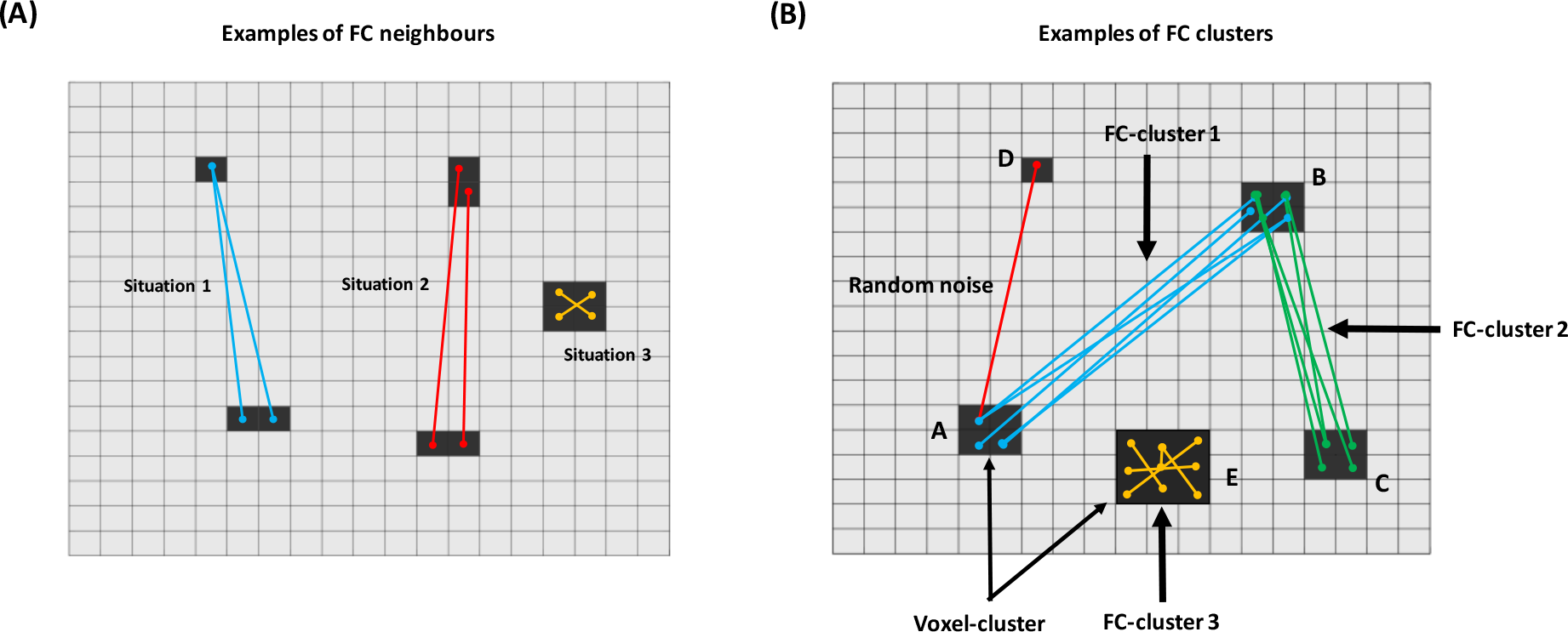
Two-dimensional diagrams of FC neighbours and FC clusters. The size of FC clusters exceeding a CDT is used as a test statistic in the cluster-level inference. (A) In BWAS, there are two cases that FCs are neighbours. In situation 1, two FCs share a common endpoint and another two endpoints are spatial neighbours. In situation 2, two pairs of endpoints of two FCs are all spatial neighbours. Situation 3 is a special case of situation 2. (B) In BWAS, FCs can form a cluster in two ways. The first one is the FCs between voxel cluster AB, BC and AD, and the second one is FCs within a voxel cluster E.

Based on the normality and stationarity assumption as in peak-level inference, we propose to use Gaussian random field theory to approximate the null distribution of maximum cluster size. In brief, let *M* be the number of FCs exceeding a pre-specified CDT *z*_0_, *N* be the number of FC clusters, and *S* be size of a FC cluster. Suppose that separate FC clusters are independent, then the distribution of maximum cluster size *S_max_* for Gaussian random field is [Adler, 1981; Friston et al., 1994]:

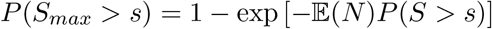

The expected number of FC clusters 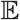(*N*) at a high CDT *z*_0_ can be approximated by the expected EC of Gaussian random field using equation (2):

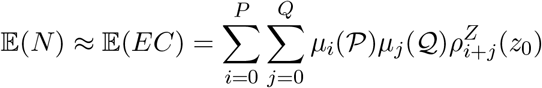

The distribution of *S* can be approximated by [Adler, 1981; Nosko, 1969]:

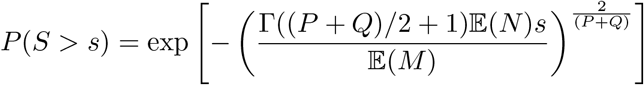

and 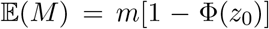, where *m* is the number of functional connectivities, and Φ(•) is the cumulative distribution function of standard normal distribution. The above theory is a generalization of previous result (e.g. [Friston et al., 1994] and [Hayasaka and Nichols, 2003]), except the use of equation (2) to approximate the expected number of clusters 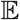(*N*).

Therefore, in the cluster-level inference, small-sized FC clusters are more likely to be identified as false positives and filtered out (e.g. the red link in Figure 3B). Large-sized FC clusters represent an existence of association signals either between two different voxel clusters (e.g. the blue and green ones in Figure 3B) or within a single voxel cluster (e.g. the yellow one in Figure 3B). For example, consider that one performed a case-control BWAS, then the identified FCs can be either altered connections between two brain regions or within a single region.

### 2.3 Validating peak- and cluster-level inference in real data

#### 2.3.1 Data

Two resting-state fMRI datasets are used in our analysis: (1) 197 subjects from the Cambridge dataset in the 1000 Functional Connectomes Project (1000 FCP); (2) 222 subjects from the Southwest University (SWU) dataset in the International Data-sharing Initiative (IDNI). The subjects in the two datasets are all healthy people with similar demographic information. They are preprocessed using standard preprocessing pipelines implemented in Data Processing and Analysis for Brain Imaging (DPABI) [Yan et al., 2016]. Finally, All fMRI data are registered to 3 × 3 × 3 mm^3^ standard space, and 47636 voxel time series within each subject’s 90 cerebrum regions (based on AAL template) are extracted. They are then smoothed by 3D Gaussian kernels with FWHM = 0, 2, 4, 6, 8, 10, 12 mm on each dimension. Therefore, for volume-based fMRI data, a total number of 14 datasets (2 sites × 7 smoothness) are used in our subsequent analysis.

In addition, the above data are also mapped on to the Conte69 surface-based atlas using the Connectome Workbench software. They are smoothed by 2D Gaussian kernels restricted on the cortical surface with FWHM = 0, 4, 8 mm. Finally, 32492 vertex time series on the left cortical surface are used in our analysis. All details are provided in the Appendix.

#### 2.3.2 Estimating the empirical FWER

To evaluate whether the random field theory can actually control the FWER in real data analysis, we compared our method with empirical permutation results in real data. Similar approaches have previously been adopted to validate the random field theory in task-activation studies [Eklund et al., 2016, 2012].

The following procedures were carried out in each of the volume-based and surface-based fMRI datasets. First, subjects were randomly divided into two groups. Second, BWAS was performed to compare the whole brain functional connectivities between two groups (approximately 1.13 × 10^9^ connections in volume-based data, and 5.28 × 10^8^ connections in surface-based data). The peak- and cluster-level inference approaches were applied to find significant signals. Third, the above two steps were repeated 2000 times. FWER was then estimated by computing that proportion of permutations in which any significant signal is found. Since subjects were all healthy people with similar demographic informations, and their group labels were randomly assigned, we expected that there were no group differences. Therefore, if the proposed approach is valid, the proportion of analysis with at least one significant effect should be close to the nominal error rate 0.05.

### 2.4 Comparing with other multiple correction methods

We compared our proposed method with connexel-wise Bonferroni correction and false discovery rate (FDR-BH [Benjamini and Hochberg, 1995], FDR-BY [Benjamini and Yekutieli, 2001]) in terms of the observed power and false-discovery rate. To mimic real data, we did not use completely simulated data, but rather, we adopted a widely used evaluation methodology in GWAS (e.g. [Yang et al., 2014; Zhou and Stephens, 2012]), which directly simulated signals that correlated with real data. The data used here were 197 subjects in the Cambridge dataset in four smoothness levels (FWHM = 0, 4, 8, 12 mm).

#### 2.4.1 Simulation procedures

In detail, two cerebrum regions within the AAL template were first randomly selected. BOLD signals of voxels within these two regions were extracted and functional connectivities of pairwise voxels between these two regions were estimated. Second, subjects were randomly divided into two groups, and signals were added to a subset of functional connectivities in one group. Specifically, the signals formed a single FC-cluster with different mean connectivity intensity between the two groups. Third, a two-sample t-test was used to compare two groups of functional connectivities. Five methods, including Bonferroni, FDR-BH [Benjamini and Hochberg, 1995], FDR-BY [Benjamini and Yekutieli, 2001], peak-level inference, and cluster-level inference (with different CDT), were used to control the false-positive rate of multiple hypothesis testings.

Four free parameters were found in our simulation: 1) voxels selected from real data, 2) signal width, i.e., the number of altered functional connectivities, 3) effect size of the signal and 4) image smoothness. In the Results Section, we report the results of comparisons among the different combinations of parameters.

#### 2.4.2 Performance metrics

Two metrics were used to evaluate the performance: the observed power and false-discovery rate. The observed power was calculated as the number of discovered true-positive functional connectivities divided by the total number of true-positive connectivities. The observed false-discovery rate was calculated as the number of discovered false-positive functional connectivities divided by total number of discovered functional connectivities.

### 2.5 Statistical power analysis

A method to estimate the statistical power of peak-level inference is proposed. Power is defined as the probability of finding at least one true-positive signal for a region (denoted as B) in which the false-positive rate α is controlled at a certain level in the whole search region (denoted as A) [Friston et al., 1994]. To estimate power, four parameters should be specified: (1) the threshold that controls FWER *α*, (2) the effect size of true signal *γ*, (3) the sample size *n*, and (4) the smoothness of image.

First, if we assume that the primary variable of interest, 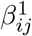, is subject to a normal distribution 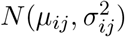, the null hypothesis is 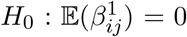. Therefore, every test statistic *Z_ij_* is subject to *N*(0,1). The whole search region *A* is a central Gaussian random field with mean zero and variance one at each point. The threshold *z*_0_ to control the FWER at *α* is obtained by the random field theory (Formula 2):

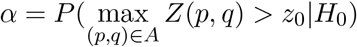

where (*p, q*) are the coordinates of the functional connectivities.

Then, under the alternative hypothesis 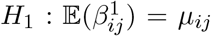, the test statistics *Z_ij_* is subject to 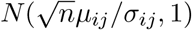, where *n* is the sample size. The *γ_ij_* = *μ_ij_*/*σ_ij_* is called effect size at FC*_ij_*. We further assume that the distribution of signals will be the same in region *B*, i.e., all 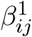 is subject to the same normal distribution *N*(*μ*, *σ*^2^). Therefore, region *B* is a non-central Gaussian random field *Z*\*(*p, q*) with mean 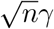 and variance one at each point. The power in the search region *B* ⊂ *A* can be expressed as:

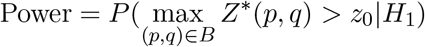

The non-central Gaussian random field *Z*\*(*p, q*) can be transformed to a central Gaussian random field by the following element-wise transformation:

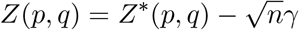

therefore, the power in region *B* can still be calculated using Formula (2):

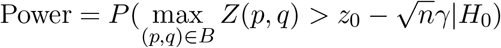

Three issues remain. The first involves selecting region *B*. When estimating power, we select region *B* as consisting of functional connectivities between two three-dimensional balls, with the diameter of each ball being equal to the intrinsic FWHM of the image (Figure 2). The idea is that signals within such ball are usually homogeneous as a result of the smoothness of the image. Besides, the matched filter theorem suggests that the signal is best detected when the width of the smooth kernel matches the width of the signal [Worsley et al., 1996a].

The second issue involves the random field theory which can only approximate the right tail of the maximum distribution. Therefore, the above method may lead to an inaccurate estimation when 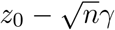 is small. To address this problem, we propose to use the following heuristic modification:

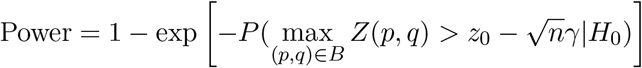

This formula ensures that the power is between zero and one, which shows excellent performance in the simulation.

The last issue concerns estimating the effect size, which is typically estimated from the statistical map of a pilot study using the same study design. Suppose that the pilot BWAS study used *n*^⋆^ samples. Then, the estimated effect size at FC_*ij*_ is [Joyce and Hayasaka, 2012]:

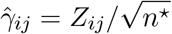

Using the above formula, the power can be estimated for each functional connectivity to form a power map on six-dimensional space, but it is quite difficult to visualize such a maps. Therefore, to report the power of a study, we estimate the effect size of every FC to form an empirical distribution. The power curves of different sample sizes and effect sizes under certain power (e.g. 90% power) are analyzed and reported.

#### 2.5.1 Simulation-based validation for power analysis

To test whether the proposed method can estimate power accurately, we performed a simulation study. Briefly, we simulated a case-control study with known effect size in a subset of functional connectivities, and we compared the observed power and the theoretical power.

In detail, first, we generated two sets of 10000 three-dimensional independent Gaussian white noise images, with 30 voxels per dimension. Second, the images were smoothed by Gaussian kernels with FWHM ranging from 3 to 6 voxels. Third, a ball with radius 10 voxels located at the center of each image was extracted. This guaranteed the uniform smoothness. Fourth, every 20 images were combined to form 500 simulated four-dimensional fMRI data. We denoted the images in the first set as (*A*_1_, *A*_2_,…, *A*_500_) and the images in the second set as (*B*_1_, *B*_2_ *B*_500_). Fifth, the Pearson correlation coefficients were calculated between time series of pairwise voxels of images *A_i_* and *B_i_*, and a Fisher’s Z transformation was then performed. Sixth, two groups of images from two sets were randomly selected, with each group consisting of 200 samples. A *Z*-map was then generated by fitting each functional connectivity to a general linear model to compare the two groups. Seventh, signals were then added to functional connectivities between two balls, which were located at the center of each images. The diameter of balls was equal to the FWHM of images. Specifically, a signal was the mean intensity difference between two groups. We then estimated power using simulated data under different parameters, including image smoothness FWHM, sample size *n* and effect size *γ* (Figure 2). The steps six and seven were repeated for 10000 times under each parameter setting, and its maximum statistics are recorded at each simulation. The empirical power was estimated by the proportion of maximum statistics exceeding the FWER 0.05 threshold. We compared the results of simulation with the proposed theoretical method.

## 3 Results

### 3.1 Overview of the proposed approaches

Figure 1 and 2 show the diagrams of the proposed approaches. Figure 4 shows the multiple comparison threshold of different approaches in a typical BWAS study. In the study, the fMRI data have a spatial resolution of 3 × 3 × 3 mm^3^. A total of 47636 voxels in the cerebrum regions were used.

**Figure 4:**
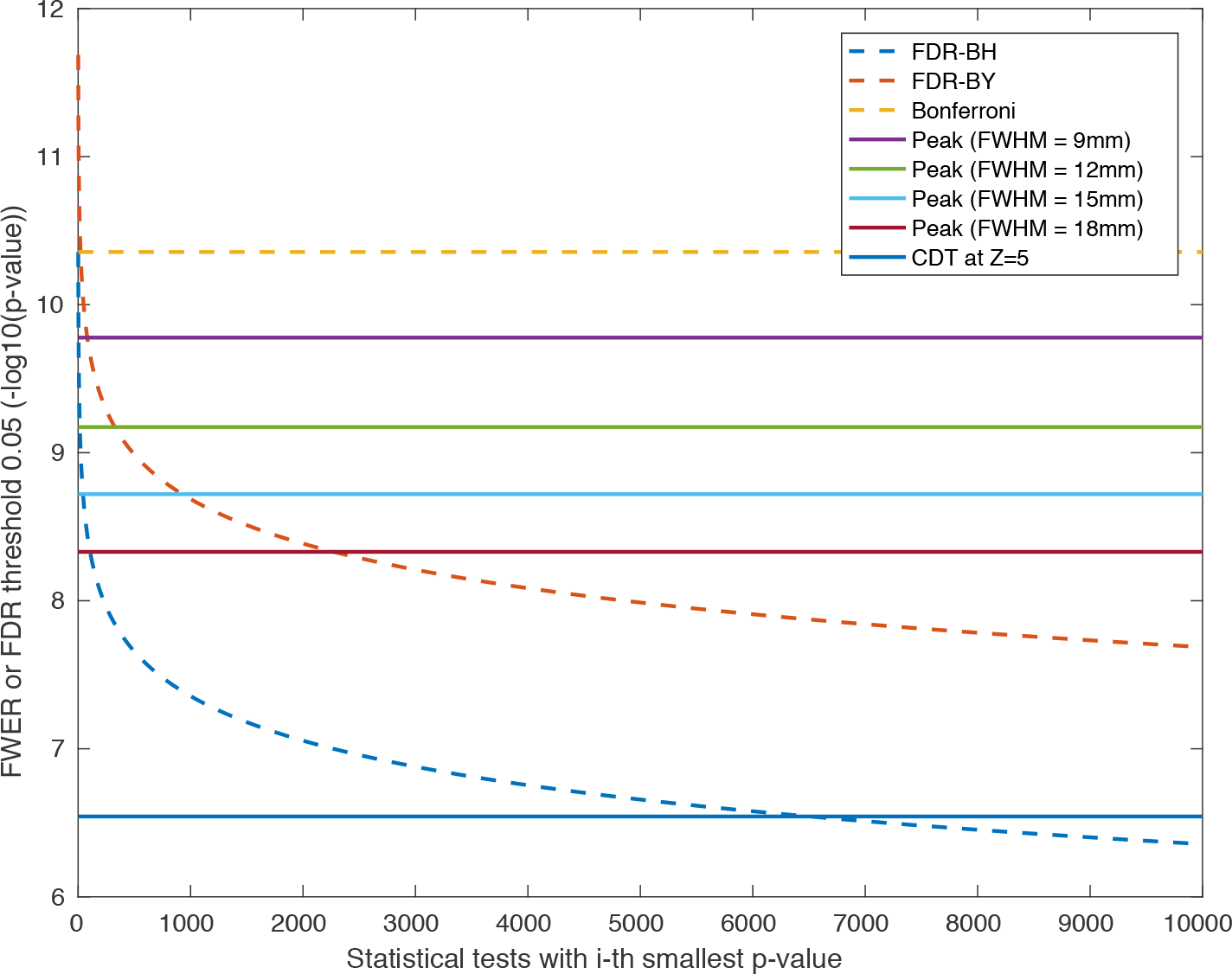
A comparison of the multiple comparison thresholds provided by different methods in BWAS (FWER or FDR 0.05) using 3×3×3 mm^3^ volume-based fMRI data. Methods that control FWER (Bonferroni correction and peak-level inference) provide universal thresholds across FCs. The threshold of FDR approaches depend on the rank of p-values of FCs. The CDT of cluster-level inference is an universal threshold across FCs, and a subsequent correction on the size of FC clusters is applied.

Methods that control connectivity-wise FWER, including Bonferroni correction and peak-level inference, provide evidence of association of each individual functional connectivities that survive the threshold. Bonferroni correction is always the most conservative one. The peak-level inference is more powerful when the smoothness of images are increased. As shown in the Figure 4, the FWER-corrected threshold can be 1 to 2 order of magnitudes less conservative than Bonferroni correction. Methods that control connectivity-wise FDR, including FDR-BH and FDR-BY approaches, control the proportion of false-positive findings smaller than a pre-specified level *q* (e.g. 5%). For the widely-used FDR-BH approach, it compares the *i*-th smallest p-value *p*_(*i*)_ with 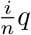, where *n* is the total number of hypothesis tests, and rejects the first *k* hypothesis tests that satisfy 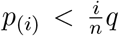(Figure 4). Therefore, the power of FDR approaches highly depends on the observed p-values, which can be more or less powerful than the peak- and cluster-level inference. For example, FDR approaches require the most significant p-value reaches the threshold of Bonferroni correction. This requirement is sometimes very conservative in BWAS, owing to the billions of statistical tests performed. However, it can be more powerful when many of the p-values meet the requirement of the data-driven threshold. A method that controls connectivity-wise FWER can also control connectivity-wise FDR. The cluster-level inference approach tests the size of the FC clusters exceeding a CDT. A significant FC cluster can provide evidence that there exist association signals somewhere in this FC cluster. None of the individual functional connectivities in the cluster can be declared as significant ones. This approach is usually sensitive to spatially extended signals. Moreover, when the CDT equals the FDR threshold, the connectivity-wise FDR can be controlled, and when the CDT equals the FWER threshold, it is equivalent to control the connectivity-wise FWER.

### 3.2 Validating peak- and cluster-level inference in real data

#### 3.2.1 Estimated FWER in real datasets

We evaluate whether the proposed method can control the FWER in real data analysis by comparing the theoretical FWER with the empirical FWER estimated by permutation approaches. The experimental procedures are illustrated in Section 2.3.2. For volume-based fMRI data, we used 14 datasets (2 sites × 7 smoothness). For surface-based fMRI data, we used 6 datasets (2 site × 3 smoothness). The estimated smoothness of different datasets are shown in Table 1.

**Table 1:** The estimated smoothness of different datasets used in our analysis (FWHM in mm).

| Applied smoothness |  | 0 mm | 2 mm | 4 mm | 6 mm | 8 mm | 10 mm | 12 mm |
| --- | --- | --- | --- | --- | --- | --- | --- | --- |
| Estimated smoothness | Cambridge volume-fMRI | 4.5 mm | 4.6 mm | 7 mm | 10.6 mm | 13.2 mm | 15.9 mm | 17.6 mm |
|  | Cambridge surface-fMRI | 4.7 mm | NaN | 6.9 mm | NaN | 11.1 mm | NaN | NaN |
|  | SWU volume-fMRI | 5.1 mm | 5.2 mm | 7.5 mm | 10.9 mm | 13.4 mm | 15.3 mm | 17.9 mm |
|  | SWU surface-fMRI | 5.2 mm | NaN | 7.7 mm | NaN | 12.2 mm | NaN | NaN |

Figure 5 shows the estimated FWER of peak- and cluster-level inference methods using volume-based fMRI data. We found that the peak-level approach is valid, as most of the estimated FWERs lie in the binomial confidence interval of 2000 permutations (dashed line). The cluster-level inference is also valid if the CDT is larger than 5. However, when the CDT becomes smaller, the false-positive rate will exceed the nominal level, because the assumptions of the theory may break down.

**Figure 5:**
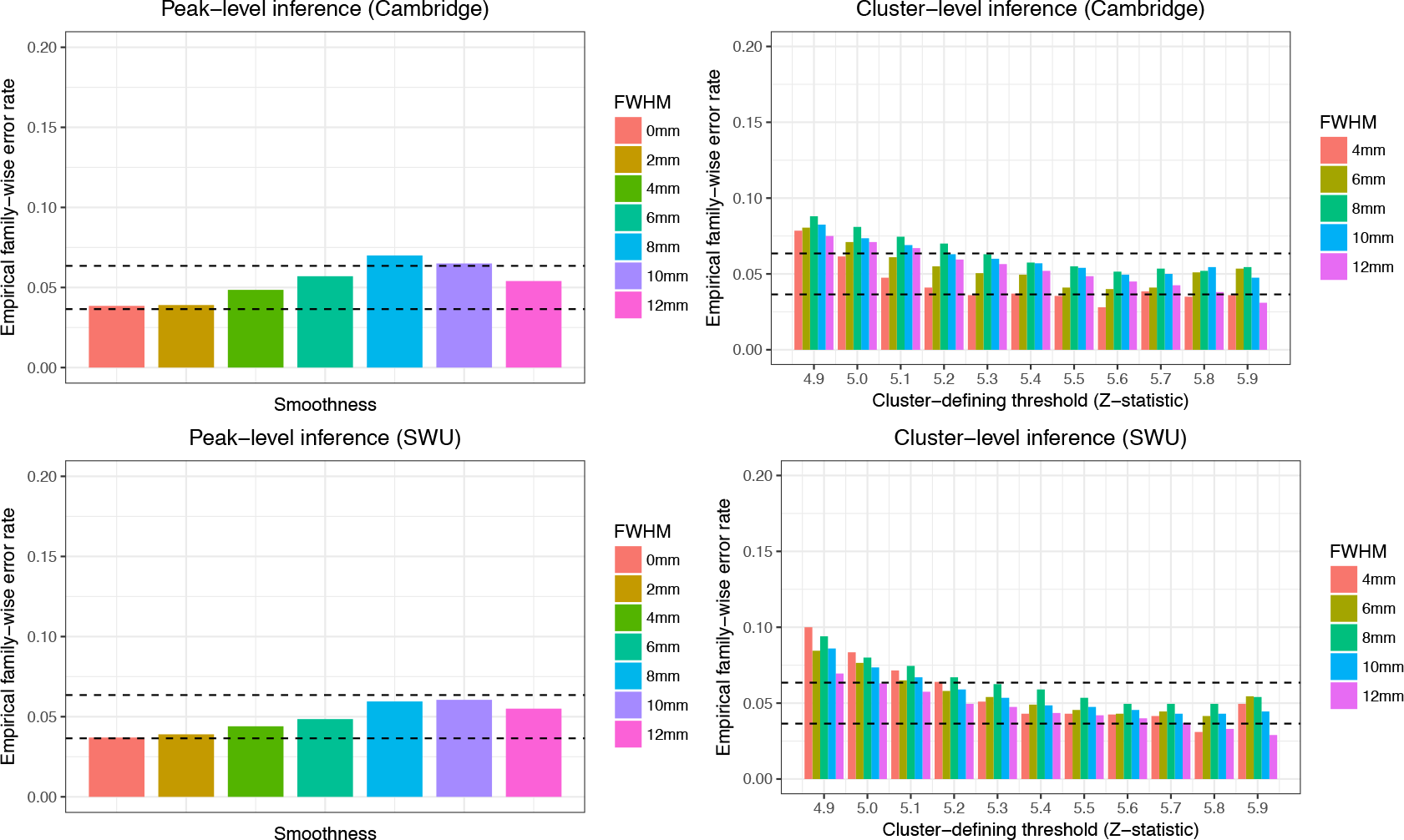
Validating peak- and cluster-level inference by comparing the theoretical FWER with permutation-based empirical FWER at 0.05. The methods are tested in 2 datasets (top: Cambridge; bottom: SWU) under 7 different smoothness levels (0 to 12 mm smoothing). The estimated FWER is that proportion of permutations in which any significant signals are found by the random field theory. Left: Results for peak-level inference. Right: Results for cluster-level inference with different CDT (from 4.9 to 5.9). Almost all the results lie in the binomial 95% confidence interval (the dashed line).

Figure 6 shows, for cluster-level inference, the comparison of the estimated cluster-size threshold of random field theory and permutation approach at low smoothness levels. Different from the above analysis, we directly compare two thresholds because the 95% quantiles of empirical maximum cluster-size distribution can not be estimated accurately. The reason is that when the smoothness is low, the size of FC clusters is usually small, thus, there are many ties in the maximum cluster-size distribution. A good agreement between the two thresholds demonstrates the validity of cluster-size inference at low smoothness level, and the CDT can even be lower comparing with the above analysis (Z=4.5).

**Figure 6:**
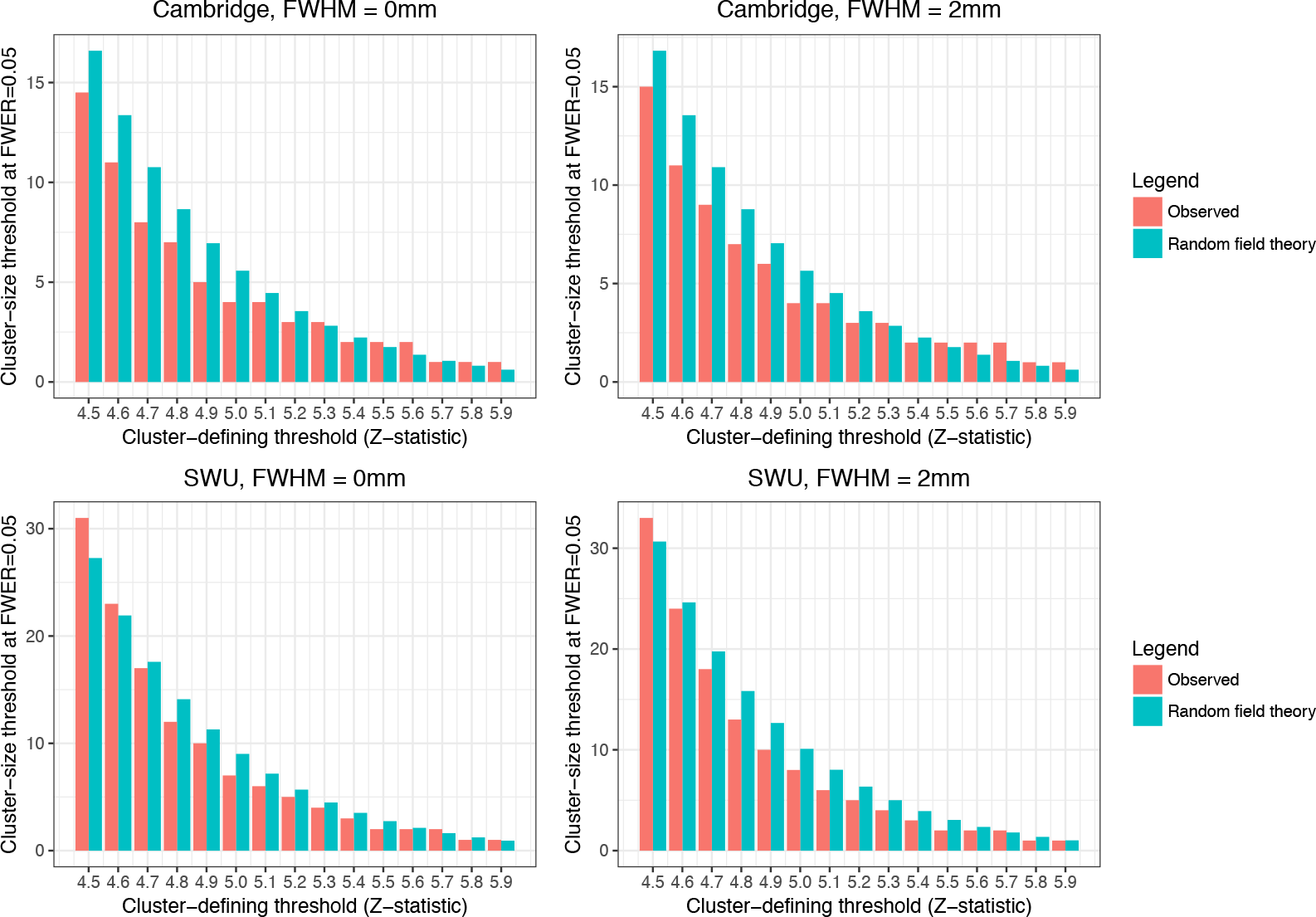
Validating cluster-level inference *at low smoothness* by comparing the theoretical cluster-size threshold with permutation-based empirical threshold at FWER 0.05 with different CDTs (from 4.5 to 5.9). The methods are tested in 2 datasets (top: Cambridge; bottom: SWU) under 2 different smoothness levels (0 mm and 2 mm smoothing).

Figure 7 shows the estimated FWER of peak- and cluster-level inference methods using surface-based fMRI data. We found that, when no spatial smoothing is applied (FWHM = 0mm), our approach is more conservative than permutation approach. The method works well when we smooth the data. However, to the best our knowledge, there are no standard preprocessing pipelines for surface-based resting-state fMRI data, thus, our surface-mapping approach may not be optimal for BWAS and different preprocessing pipelines may affect the performance of our approach. Therefore, the robustness of the approach should be tested in the future.

**Figure 7:**
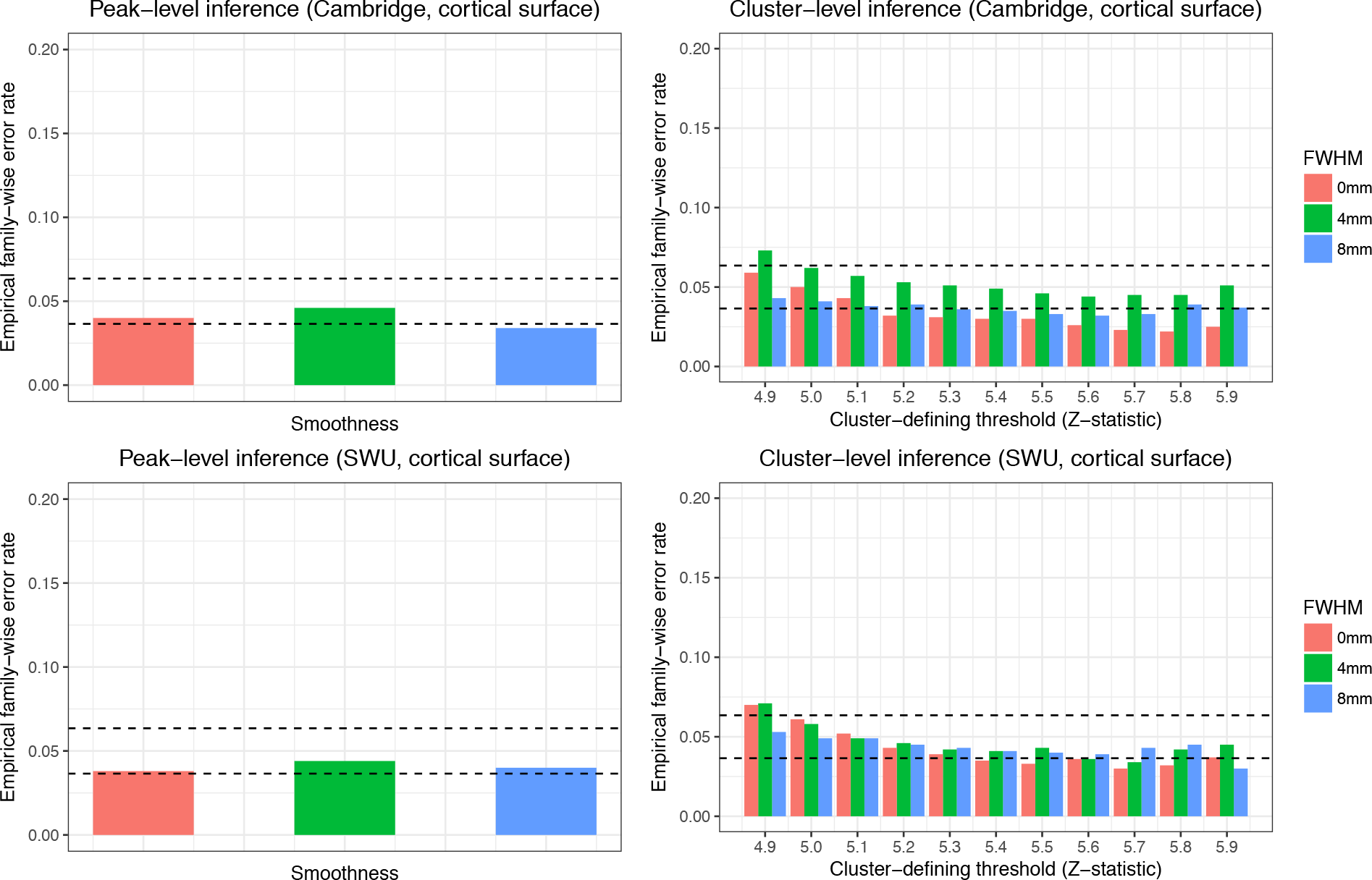
Validating peak- and cluster-level inference in *surface-based fMRI data* by comparing the theoretical FWER with permutation-based empirical FWER at 0.05. The methods are tested in 2 datasets (top: Cambridge; bottom: SWU ) under 3 different smoothness levels (0, 4, 8mm smoothing). The estimated FWER is that proportion of permutations in which any significant signals are found by the random field theory. Left: Results for peak-level inference. Right: Results for cluster-level inference with different CDTs (from 4.9 to 5.9). Almost all the results lie in the binomial 95% confidence interval (the dashed line).

#### 3.2.2 The choice of cluster-defining threshold

For cluster-level inference, the expected Euler Characteristics is used to approximate the expected number of clusters in the random field theory, assuming that the absence of holes when CDT is applied. However, this assumption may not be true when the CDT is not high enough or the data are not smooth enough. Therefore, we compare the expected Euler characteristics calculated based on the Gaussian random field theory with the observed expected number of clusters across different levels of CDT in volume-based fMRI data in both Cambridge and Southwest University datasets. The observed expected number of clusters is computed based on an average of 2000 permutations of each dataset. The results are shown in Supplement Figure 2 and 4. We found that, when the applied smoothness is larger than 4mm, the choice of CDT greater than 5 is very safe for 3 × 3 × 3 mm^3^ resolution fMRI data to meet the assumption of the random field theory. This is in agreement with our results in the previous Section (Figure 5). When the smoothness is low, we found that there exist a large deviation between theory and real data when the CDT is smaller than 5.5. However, the results shown in Figure 6 indicate the proposed method can provide a valid threshold when the CDT is as low as 4.5 in two datasets. Therefore, more analysis should be done to validate the approach in the low smoothness cases.

#### 3.2.3 Distribution of functional connectivity data

We test whether functional connectivities data, i.e., Fisher’s Z transformed correlation coefficients, are subject to normal distributions, which is a critical assumption for Gaussian random field theory. We performed one-sample Kolmogorov-Smirnov test to test the normality of each functional connectivity in both Cambridge and Southwest University datasets. Supplement Figure 1 and 3 show the results. As most of the p-values are larger than 0.05, we conclude that the normality assumption is met.

### 3.3 Comparing peak- and cluster-level inference with other multiple correction methods

Figure 8 shows the results of comparisons using 197 subjects in the Cambridge dataset. The experimental procedures are illustrated in Section 2.4. In this analysis, we extracted time series of 306 voxels from the left putamen region and 302 voxels from the left inferior frontal gyrus in each of the 197 subjects, and 306 × 302 = 92412 functional connectivities between these two regions were calculated. Signals were added to 2970 of the functional connectivities, with effect size ranging from 0.15 to 0.3, and smoothness of 0, 4, 8, 12 mm was applied. For cluster-level inference, we used CDT = 3, 3.5 and 4 (Z-value). The following observations are obtained from this simulation:

**Figure 8:**
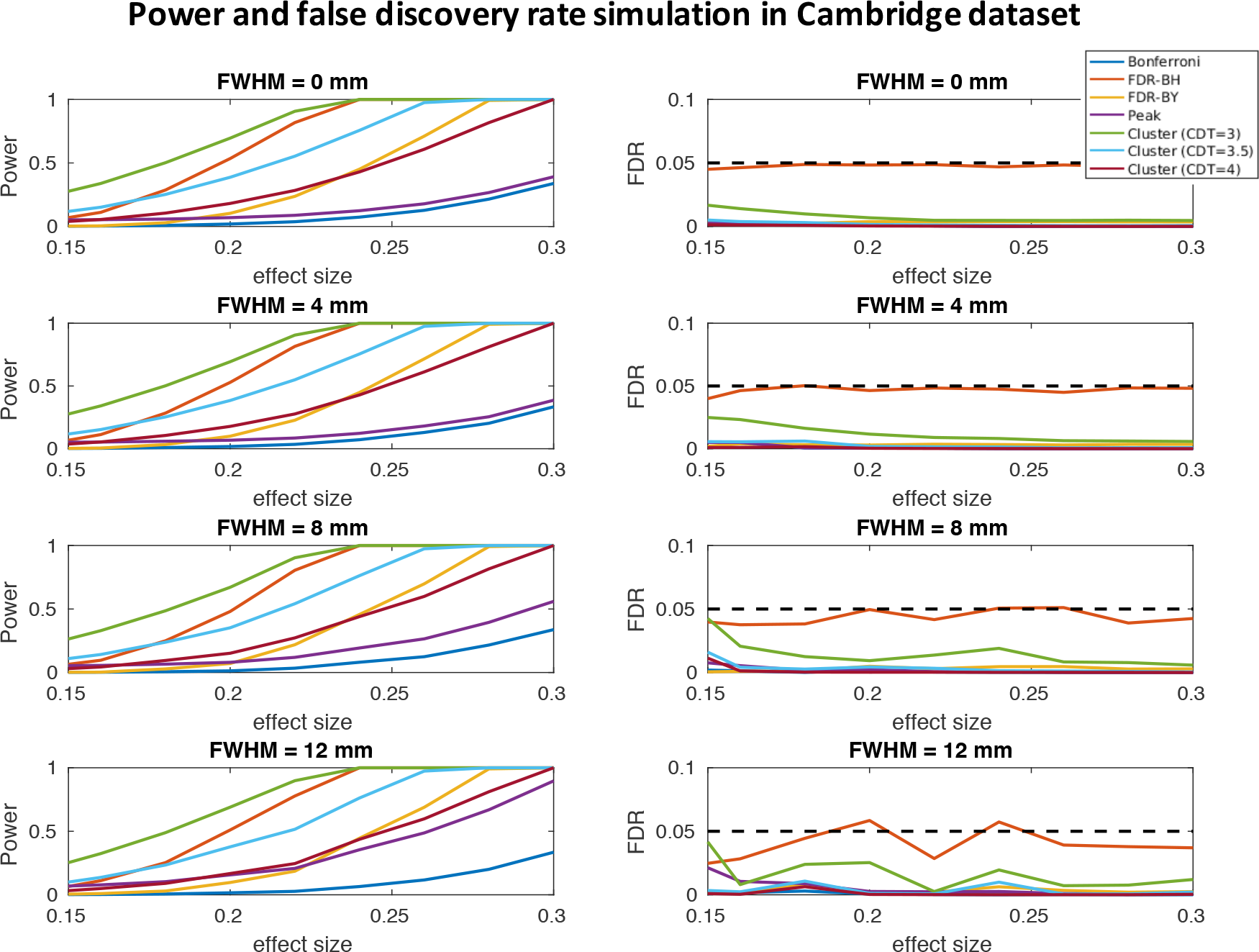
Comparing peak- and cluster-level inference methods with Bonferroni correction and two FDR methods in terms of power (first column) and false discovery rate (second column) across different levels of effect size and smoothness levels using the Cambridge dataset. Left: power curves of different approaches under different smoothness levels. Right: FDR curves of different approaches under different smoothness levels.

- Almost all the methods can control false discovery rate in this simulation (below 5%).
- The power of peak-level inference is similar to Bonferroni correction when the smoothness is low (e.g. no spatial smoothing), but it becomes close to FDR-BY and much higher than that of Bonferroni correction when the smoothness is high (e.g. applied smoothness of 12 mm).
- The false discovery rate and power of cluster-level inference depends on the choice of CDT. The lower the CDT, the higher the false discovery rate and power.
- We can find a CDT whose power is higher than that of the FDR method. In the meantime, the false discovery rate is lower. For example, for cluster-level inference with CDT=3, the power is higher than that of the FDR-BH method, and the false discovery rate is lower. For cluster-level inference with CDT=3.5, the power is higher than that of the FDR-BY method, and the false discovery rate is lower.

Similar results can be obtained by changing the selected voxels and the width of the signal added, as shown in the Appendix (Supplement Figure 5-7). In conclusion, cluster-level inference can increase sensitivity and decrease false-positive rate by filtering out small FC-clusters generated by random noises. Peak-level inference shows increased power when the smoothness is large; thus, it is recommended when performing group-level studies with large applied smoothness.

### 3.4 Real data analysis: identifying altered functional connectivities in major depression disorder

We applied our method to identify functional connectivity difference between patients with major depression disorder (MDD) and healthy controls. The data used here are part of the data in our previous study [Cheng et al., 2016] which contained 282 patients and 254 demographic information matched controls from Southwest University dataset. We applied BWAS approach to test the connectivity difference between two groups, with age, gender, education year, head motion (mean frame-wise displacement) being nuisance covariates.

The most significant p-value among all functional connectivities was *p* = 5.5 × 10^−11^. However, the Bonferroni correction, FDR-BH and FDR-BY approaches can not detect any significant connectivities (FWER or FDR at 0.05). This is because Bonferroni correction requires the p-value smaller than *p* = 4.4 × 10^−11^, and both FDR-BH and FDR-BY approaches require the most significant p-value smaller than the same threshold as Bonferroni correction. See the Manhattan plot for details (Figure 9).

**Figure 9:**
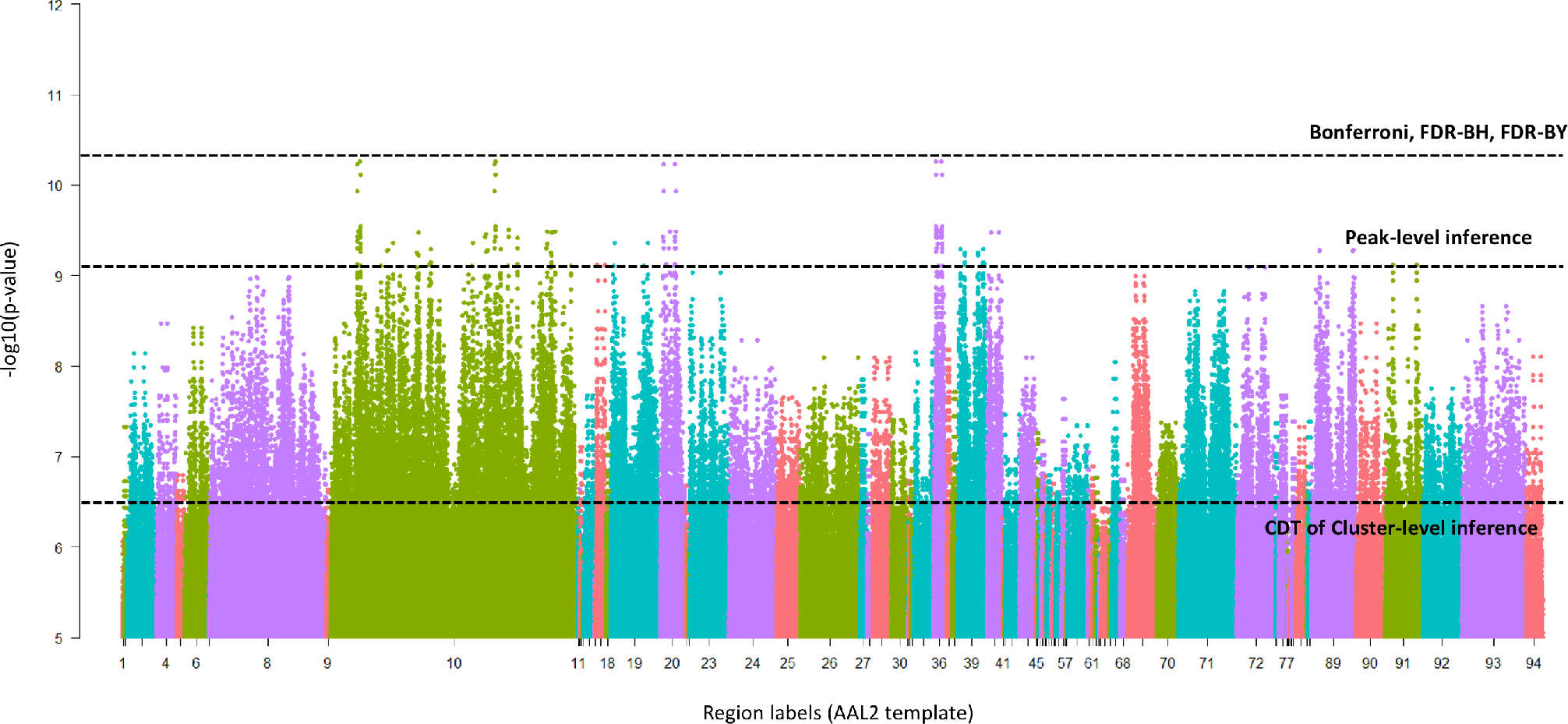
Manhattan plot of altered functional connectivities in major depression disorder (p < 10^−5^ only). Each point represents a functional connectivity grouped by the 94 cerebrum regions of the AAL2 template. Bonferroni correction, FDR-BH and FDR-BY fail to identify any significant connections, while both peak- and cluster-level inference approaches identified many altered connectivities. Their brain locations are shown in the next two figures.

The p-value threshold of peak-level inference approach was *p* = 9.1 × 10^−10^ (connectivity-wise FWER=0.05). A total of 114 altered functional connectivities were found (Figure 10, left). We applied cluster-level inference approach to identify significant FC clusters (CDT *p* = 3 × 10^−7^ (Z=5) and cluster-size FWER=0.05 ). A total of 12388 functional connectivities were found with p-value smaller than the applied CDT, and they formed 117 FC clusters. The largest one contains 2247 functional connectivities. Finally, 10 largest FC clusters survived the cluster-size FWER 0.05 threshold (Figure 10, right). Almost all the significant FCs in peak-level inference form FC clusters in the cluster-level inference. We could see that, although billions of hypothesis tests were performed and tens of thousands of functional connectivities were found, the results obtained by the cluster-level inference are very structured, thus, easy to be reported (Figure 10, right). The identified FC clusters can be used in subsequent analysis in several ways. For example, we can calculate the mean functional connectivity within each FC clusters, and use prediction models to classify patients and controls in a new dataset. For patients, we can also test whether the mean functional connectivity within each FC clusters are associated with the depression symptom severity scores.

**Figure 10:**
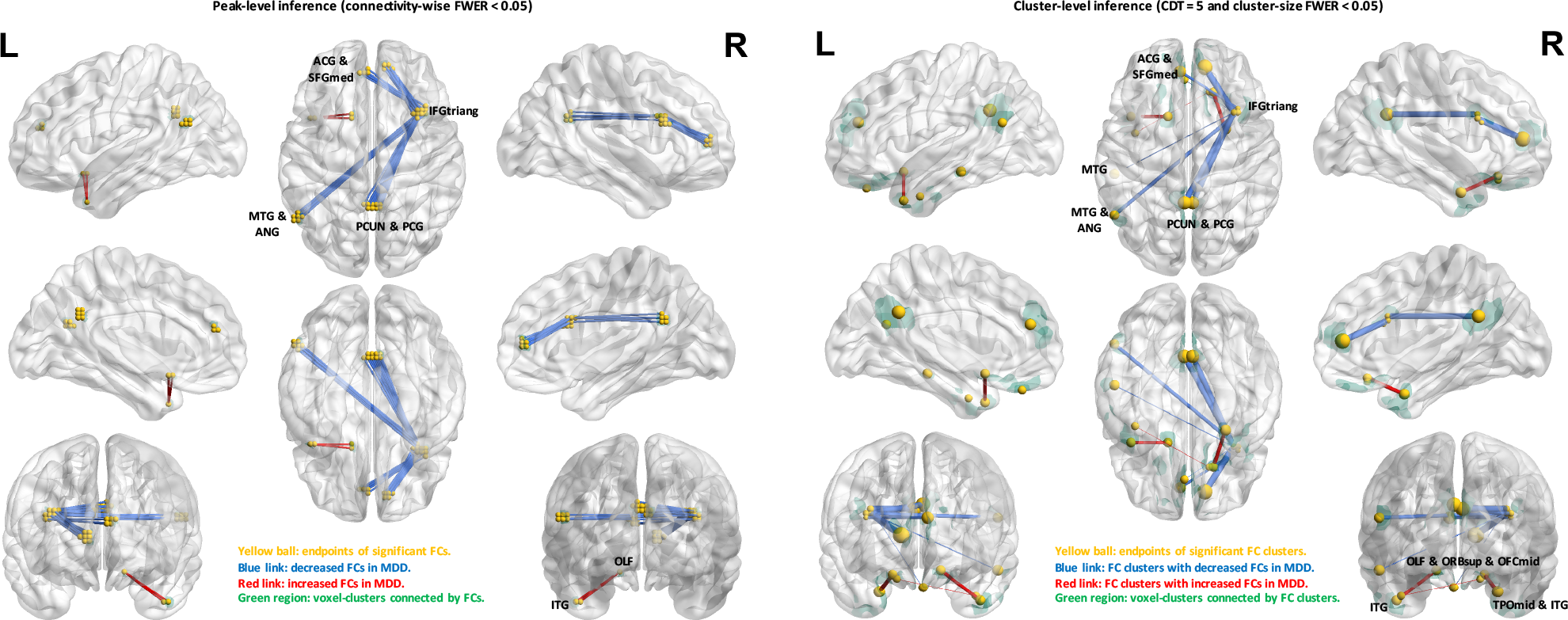
The altered functional connectivities in major depression disorder identified by peaklevel inference (left) and cluster-level inference (right). For peak-level inference, the connectivitywise FWER is 0.05, which corresponds to uncorrected p-value < 9 × 10^−10^. For cluster-level inference, the CDT is Z=5 (*p* < 3 × 10^7^) and cluster-size FWER is 0.05. Abbreviations of regions are listed in the Supplement Table 1.

### 3.5 Simulation-based validation for power analysis

Figure 11 shows the relationship between sample size and power estimated by two methods under two combination of parameters: effect size γ = 0.2, 0.4, 0.6 and smoothness FWHM=3,4,5,6 voxels. The estimation error (mean squared error) shown in the figure is very low. Therefore, the proposed method can estimate power accurately, and this proposed framework can save a considerable amount of time in generating power curves.

**Figure 11:**
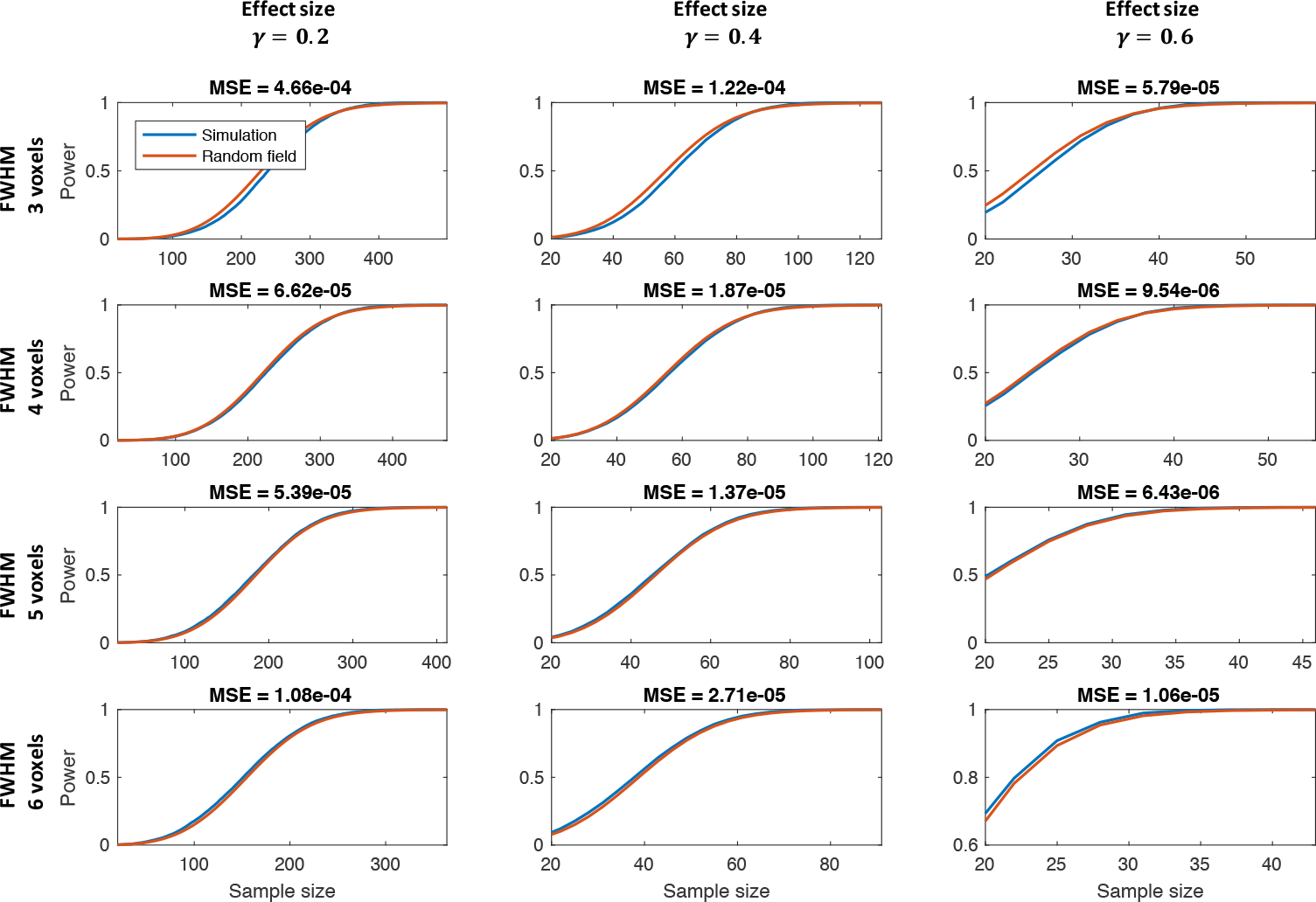
Comparing the theoretical power analysis method (red line) with the simulation result (blue line). Each figure shows the relationship between estimated power and sample size. From the left to the right, the effect sizes are 0.2, 0.4 and 0.6. From the top to the bottom, the FWHMs are 3 to 6 voxels.

### 3.6 The power of a future brain-wide association study on MDD

We show an example of how to perform a power analysis to estimate the minimum required sample size for a BWAS. In this example, we will analyze the power of BWAS on MDD using the results of the above study. The aim is to estimate the minimum required sample size to find at least one altered functional connectivities. Base on the above study, the most significant functional connectivities is *p* = 5.5 × 10^−11^, corresponding to an effect size of *γ* = 0.28.Assuming that in the new dataset, this functional connectivity has a similar effect size, the power under different sample sizes and smoothness levels are estimated and plotted in the Figure 12. Results show that about 80 to 130 subjects are needed to reach 90% power under different smoothness levels.

**Figure 12:**
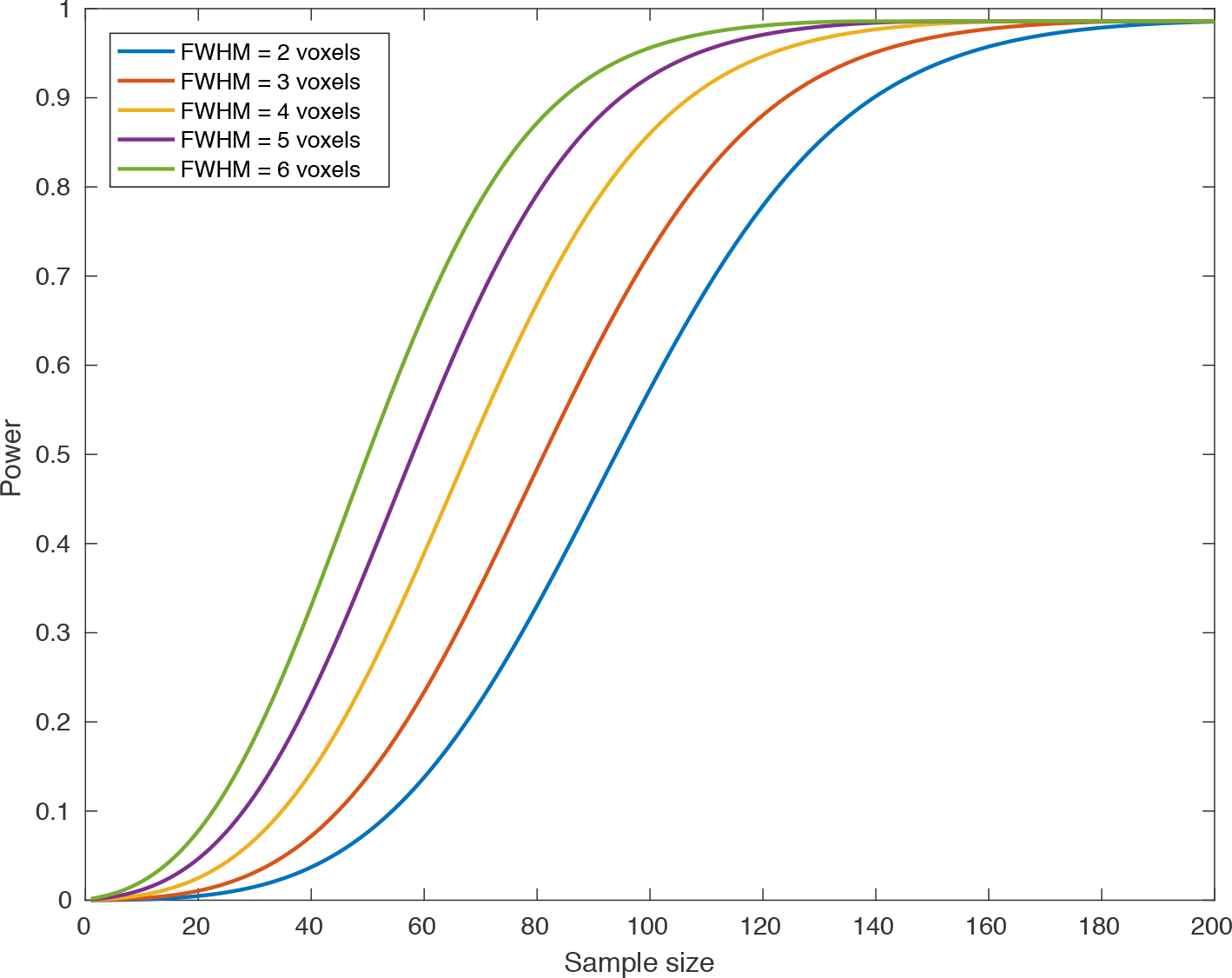
Power of detecting at least one altered functional connectivity in major depression disorder under different sample sizes and smoothness levels estimated by the proposed approach.

## 4 Discussion

Our proposed method can accurately control FWER, as demonstrated by comparing with the empirical FWER obtained from two real datasets. To the best our knowledge, BWAS is the first method to use the random field theory to analyze the voxel-wise functional connectome. Random field theory makes some assumptions of data. Eklund et al. [2016] recently reported that random field theory could lead to inflated false-positive rate in task-activation analysis, particularly when the CDT is low (*p* =0.01). This failure is well known since the choice of low CDT violates the assumptions of the original theory [Friston et al., 1994]. However, in this same article [Eklund et al., 2016], FWER is closer to nominal level when CDT is higher (*p* =0.001). Another article [Flandin and Friston, 2016] has also pointed out that the random field theory can provide acceptable FWER when using two-sample t-test instead of one-sample t-test and resampling the data close to the original image resolution. In our analysis, we have demonstrated that the random field theory is valid for both volume-and surface-based resting-state fMRI data under different smoothness. Particularly, the CDT in cluster-level inference should be high enough (|*Z*| >5 for moderate or large smoothness and (|*Z*| > 4.5 for low smoothness).

Importantly, our method is computationally efficient. It is a fully parametric approach which is not based on any simulation or permutation. Although non-parametric approaches can also perform multiple correction and power analysis [Nichols and Holmes, 2002], they are extremely slow in connexel-wise analysis by the necessity of calculating billions of statistical tests many times. Our empirical studies show that our approach is usually at least N times faster than the non-parametric permutation approaches, where N is the number of permutations performed. The reason is that, although the subject-level brain network can be computed only once in non-parametric permutations, the fitting of connexel-wise GLM is usually much slower than network construction, thus, it dominates the computation time. In addition, parallelization of permutations will not save much time, because the transmission speed of large data between processors is very slow.

There are also limitations in the current framework. The functional connectivities identified by massive univariate statistical tests approach may not be predictive, e.g., in a case-control study, the identified connectivities may not be able to classify patients and controls. A directly construction of connexel-wise prediction model is also not practical, since the model constructed on a few hundred subjects and billion of features usually has a large variance. Meanwhile, the optimization of model parameters become very difficult in this ultra-high dimensional feature space. One possible way to solve this problem is to adopt the current BWAS framework into sure independence screening (SIS) approach [Fan and Lv, 2008; Fan et al., 2009, 2010]. In SIS, each feature *x_i_*, *i* = 1,2,…, *p* is ranked in a descending order according to its correlation with the target variable *y*, and a prediction model is fitted using a subset of features whose rank is high enough. The authors showed that this intuitive approach possesses a good sure independence screening property. BWAS is a special case of the first step of SIS, thus, it is easy to be incorporate into the SIS approach. Moreover, by filtering out small-sized FC clusters using cluster-level inference approach, we expect that the prediction performance can be improved. Therefore, based on SIS, we can try to establish a connection between BWAS and prediction analysis.

Many possible extensions and improvements of the current framework can be developed in the future. First, this framework can be extended to task fMRI analysis to identify network configuration changes (e.g. [Lohmann et al., 2016]). It can support either single subject analysis or group analysis provided that the task experiment is in block design and the length of each trial is long enough to enable network construction. Second, the cluster-level inference proposed here controls the FWER of cluster size. An alternative method of controlling the FDR of cluster size was proposed in task-activation studies [Chumbley and Friston, 2009; Chumbley et al., 2010], which can be easily adopted here. Third, the estimation of subject-level functional network is based on the Pearson correlation between pairwise BOLD signal time series in the current framework, which may be suboptimal [Westfall and Yarkoni, 2016; Bellec et al., 2008; Sahib et al., 2016]. Therefore, a better approach for constructing a functional network at the voxel level should be designed and validated in the future [Narayan and Allen, 2016; Bickel and Levina, 2008]. Fourth, with the higher volume of available data, statistical methods for combining BWAS results from multiple imaging centers are needed. In BWAS, integrating results from different datasets has been shown to greatly reduce the false-positive rate and increase sensitivity [Cheng et al., 2015a, b, 2016]. However, the sample heterogeneity introduced by different sources, such as different data acquisition pipelines, population stratification, and genetic background, may make the traditional meta-analysis method used in our previous studies suboptimal.

In this paper, we developed a rigorous statistical framework for BWAS. Both peak- and cluster-level inferences are introduced for the analysis of voxel-wise functional connectomes, and the random field theory is developed to control FWER and estimate statistical power. We believe that this method will be very useful for the neuroimaging fields in the context of understanding the brain connectome.

## 5 Acknowledgments

Jianfeng Feng is supported by the National High Technology Research and Development Program of China (No. 2015AA020507), the Key Program of National Natural Science Foundation of China (No. 91230201), International (Regional) Collaborative and Exchange Program of National Natural Science (No. 71661167002), the Key Project of Shanghai Science and Technology Innovation Plan (No. 15JC1400101), and the Shanghai Soft Science Research Program (No. 15692106604), and the National Centre for Mathematics and Interdisciplinary Sciences (NCMIS) of the Chinese Academy of Sciences. Jianfeng Feng is a Royal Society Wolfson Research Merit Award holder. Lin Wan is supported by the NSFC grants (No. 11571349 and No. 11201460), the NCMIS of the CAS, the Youth Innovation Promotion Association of the CAS, and the Strategic Priority Research Program of the CAS (XDB13040600).

## Appendix of “Statistical testing and power analysis for brain-wide association study”

### Contents

A Image acquisition and preprocessing
B Calculating the intrinsic volume and Gaussian EC-density
C Estimating the smoothness of the fMRI images
D Proof of formula (1) in the main text
E Proof of formula (2) in the main text
F Supplement figures
G Supplement tables

### A Image acquisition and preprocessing

Only publicly available data are used in this article. Resting-state fMRI data are collected from two naging sites: (1) 197 samples from the Cambridge dataset in 1000 Functional Connectomes Project L000 FCP) [Biswal et al., 2010](http://fcon_1000.projects.nitrc.org/fcpClassic/FcpTable.tml); (2) 552 subjects from the Southwest University dataset in International Data-sharing Initiative DNI)(http://fcon_1000.projects.nitrc.org/indi/retro/southwestuni_qiu_index.html). All subjects are normal people. As Southwest University dataset is a longitudinal dataset, only subjects who scanned at the first time are used in this paper.

All the data collected are subject to their local ethics review boards, the experiments and the dismination of the anonymized data are approved. The detailed data acquisition methods may be found in the respective websites and papers. The data were preprocessed using SPM12 [Penny et al., 2011] tid Data Processing and Analysis for Brain Imaging (DPABI) [Yan et al., 2016]. For each individual, the preprocessing steps included discarding the first 10 time points, slice timing correction, motion correction, coregistering the functional image to individual T1 structure image, segmenting structure images and DARTEL registration [Ashburner, 2007], regressing out nuisance covariates including 24 head motion parameters [Friston et al., 1996], white matter signals, cerebrospinal fluid signals, tempo-flfiltering (0.01-0.1 Hz), normalizing to standard space of voxel size 3 × 3 × 3 mm^3^ by DARTEL, and smoothing by a 3D Gaussian kernel with FWHM = 0, 2, 4, 6, 8, 10, 12 mm. Finally, all the images are manually checked by experts to ensure preprocessing quality. Images that are not successfully preprocessed are discarded in our analysis.

The surface-based fMRI data are preprocessed using Connectome workbench. For the each volume-ased fMRI data in the Cambridge and Southwest University dataset, we map it to the Conte69 surface-ased atlas (http://brainvis.wustl.edu/wiki/index.php//Caret:Atlases/Conte69_Atlas)using the command ‘wb_command-volume-to-surface-mapping’. Each fMRI images are then smoothed by a 2D Gaus-an kernel with FWHM=0,4,8 mm using the command ‘wb_command cifti-smoothing’. Finally, the noothness of each image is estimated by the command ‘wb_command-cifti-estimate-fwhm’. The irface area of Conte69 is estimated using the command ‘wb_command-surface-vertex-areas’, which used in the random field theory.

### B Calculating the intrinsic volume and Gaussian EC-density

To perform peak-level and cluster-level inference, we should calculate the 0-to 3-dimensional intrinsic volume and the 0-to 6-dimensional EC-densities for the Gaussian random field.

Let *P* be the number of voxels, *E_x_* (or *E_y_, E_z_*) be number of *x* (or *y*, *z*)-direction edges (two adjacent voxels), *F_xy_* (or *F_yz_*, *F_xz_*) be number of *xy* (or *yz*, *xz*)-direction surface (four adjacent voxels), and *C* be the number of cubes (eight adjacent voxels). The *r_x_* (or *r_y_*, *r_z_*) be the resel size of *x* (or *y, z*)-direction, which is defined as the voxel size divided by FWHM (in mm). The 0 to 3 dimensional intrinsic volume of *S* can be calculated as:

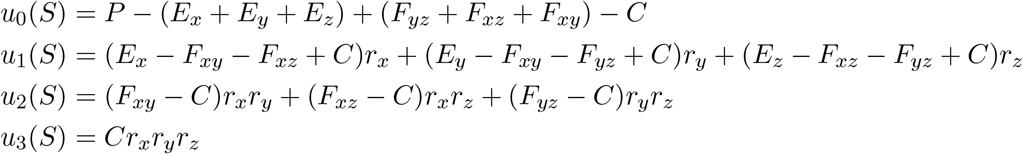

The above calculation has been implement in SPM package as *spm_resels_vol* function. Two other methods also work well in practice. One is to replace the original space with a equal volume ball, 3 implement in the *fmristat* package, the other is to use a linear regression model [Bartz et al., 2011], which do not need the knowledge of spatial smoothness. In whole-brain BWAS, for peak-level inference, the *u_i_*(𝒫) and *u_i_*(𝒬) are the same. As there are *p*(*p*-1)/2 functional connectivities across *p* axels, we divided the estimated intrinsic volume by 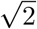, thus, the highest order term, *u*_3_(𝒫) × *u*_3_(𝒬), will approximate the total number of functional connectivities (in resel) in the brain.

The 0-to 6-dimensional EC-densities for Gaussian random field at *t* are:

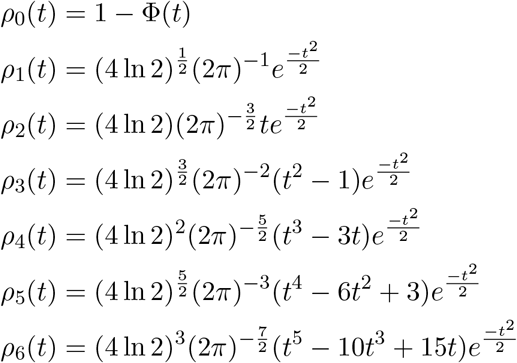

where Φ(•) is the cumulative distribution function of standard normal distribution.

### C Estimating the smoothness of the fMRI images

The true smoothness of fMRI images is usually large than the applied smoothness. Therefore, an xurate estimation of smoothness is critical for Gaussian random field theory. The following approach used to estimate the smoothness of 3D or 2D images [Hagler et al., 2006]:

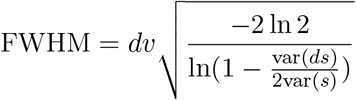

where *dv* is the average inter-neighbour distance of voxels or vertices, var(*ds*) is the variance of inter-eighbours differences, and var(*s*) is the overall variance of the values at each voxels or vertices. The FWHM of fMRI image is the average smoothness of the 3D or 2D images across all time points.

### D Proof of formula (1) in the main text

Using the property of d-dimensional intrinsic volume [Taylor and Worsley, 2008]

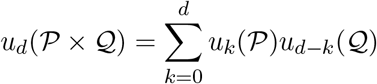

when *d* > *P, u_d_*(𝒫) = 0 and *d* > *Q*, *u_d_*(𝒬)=0. It is easy to conclude that

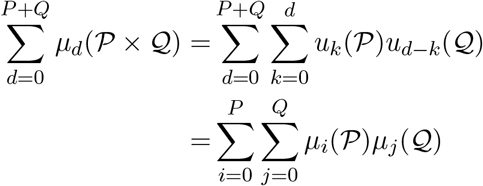

In our case, we have *P* = *Q* = 3.

### E Proof of formula (2) in the main text

The normal transformation of T-statistic makes the 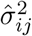 fixed as 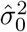 [Worsley et al., 1992, 1996]. Therere, the test statistic becomes

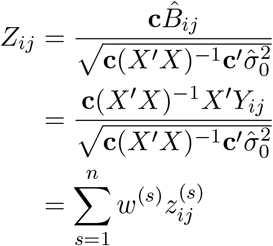

where *w*^(*s*)^ is the *s*-th element of row vector 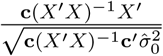, which only depends on the subjects.

Let 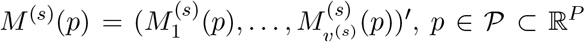 and 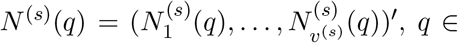 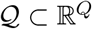 be two vectors of *v^(s)^* independent and homogeneous Gaussian random fields with mean zeros and variance one. The index *s* denotes subjects, and the *v*^(*s*)^ can be treated as the number of time oints, while *p, q* are the coordinates of three-dimensional Euclidean space. The (P+Q)-dimensional cross-correlation random field *R*^(*s*)^(*p,q*) is defined as follows [Cao et al., 1999]:

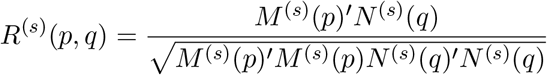

In BWAS, the cross-correlation field is generated by calculating sample correlation coefficients between pairwise voxel time series. Next, the element-wise Fisher’s Z transformation transforms this cross-)rrelation random field to a six-dimensional ‘Gaussianized’ random field as:

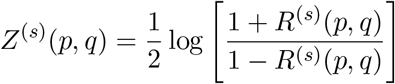

It has mean zero and variance 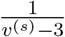 [Kenney, 1939]. Our test statistic *Z_ij_*(*p, q*) forms a weighted sum of Fisher’s Z transformed cross-correlation random field *Z*(*p, q*) as:

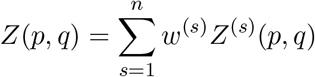

The random field *Z*(*p, q*) is a ‘Gaussianized’ random field with mean zero and variance one. Therefore, we can use formula (1) in the main text to approximate its maximum distribution at high threshold:

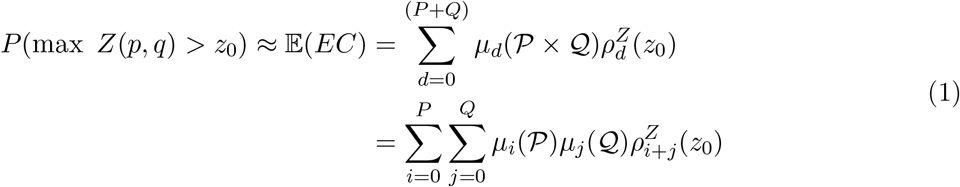

The EC-densities for the Gaussian random field 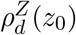 in any dimensions can be expressed as [Adler and Taylor, 2009]:

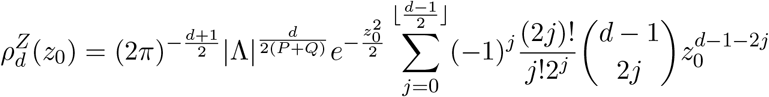

where *D* is the highest dimension of *Z*(*p,q*). The 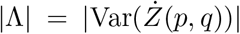 is the determinant of the variance-covariance matrix of the partial derivative of *Z*(*p, q*). The |Λ| can be replaced by FWHM_*Z*_, the Full Width at Half Maximum (FWHM) of the random field *Z* averaged across all dimensions, using the equation:

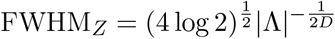

and FWHM_*Z*_ is a corrected smoothness parameter, which can be calculated as:

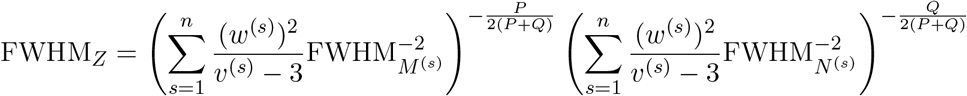

where FWHM_*M*^(*s*)^_ and FWHM_*N*^(*s*)^_ are the average FWHM of the random field vectors *M^(s)^(p)* and *N^(s)^(q)* across three dimensions. The proof of this formula is given in the next section. Finally, the formula (2) in the main text is used in the peak-level inference:

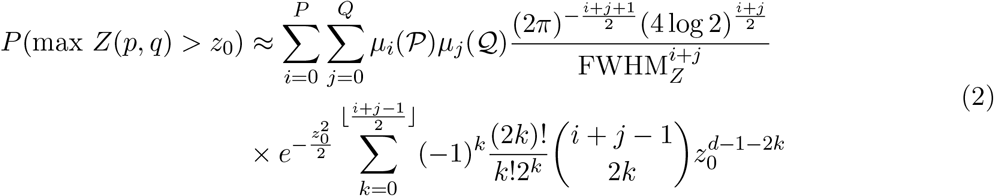

We can see that FWHM_*Z*_ is a function of the number of time points *v^(s)^* and the FWHM_*M*^(*s*)^_ and FWHM_*N*^(*s*)^_ of the individual fMRI data. Assuming that the length of scanning time and image smoothness is the same for every subject in a study. Denoting them as *v* and FWHM, the formula for calculating FWHM_*Z*_ reduces to:

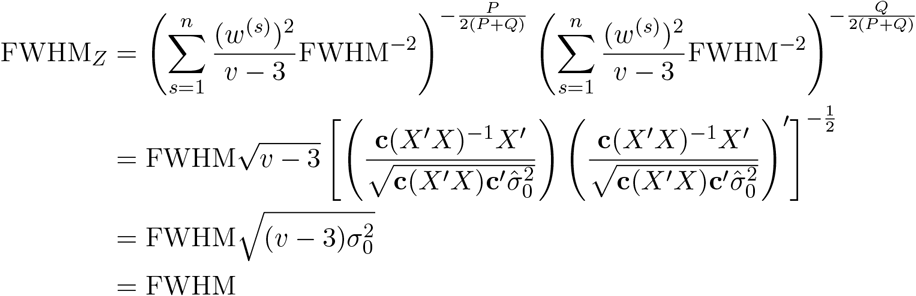

where we treat the sample variance 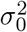 as the theoretical variance 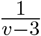. This suggests that (1) the noothness of random field *Z*(*p, q*) equals the original image smoothness. A series of non-linear trans-irmations will maintain the original smoothness of images. (2) the scanning time does not influence is formula.

In practice, for volume-base BWAS, we found that the formula one usually provide a slightly conservative estimation of FWER-corrected threshold. Therefore, we modify it as:

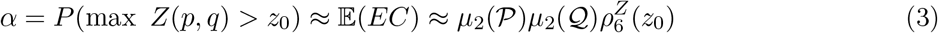

For surface-based BWAS, we use:

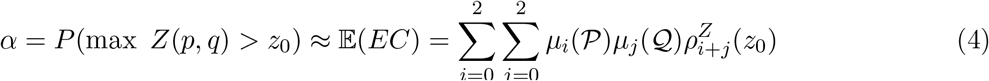

Finally, if we only analyse the functional connectivities between subcortical voxels 𝒫 and cortical vertices 𝒬, we use:

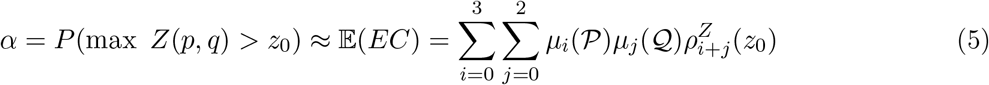

### Proof of the formula for calculating FWHM_Z_

Let 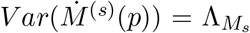 and 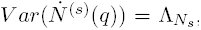, then according to the Lemma 4.2 in [Cao et al., 1999],

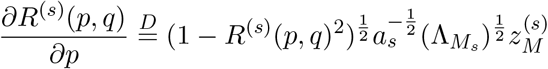

and

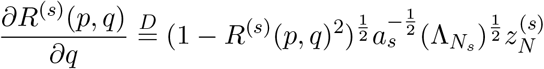

where 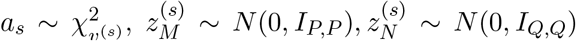 and independent of *R*^(*s*)^(*p,q*), and 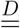 means equal in distribution. Then, after the Fisher’s Z transformation, we have

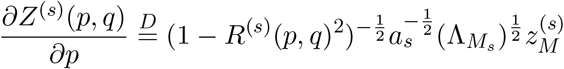

and

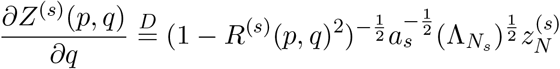

Then,

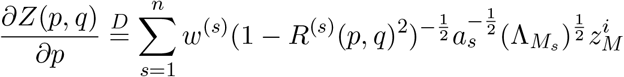

and

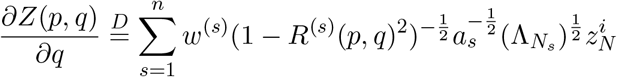

Since

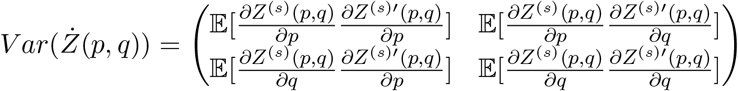

and

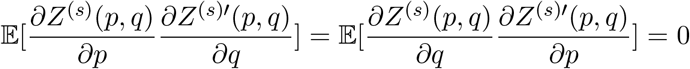

and

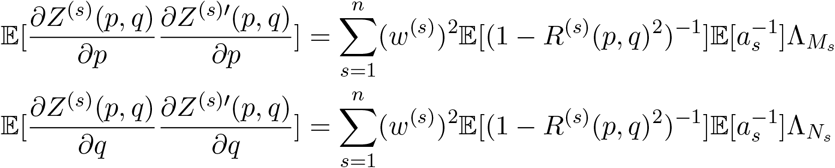

The expectations in the above equations are

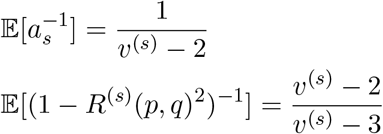

Finally we get

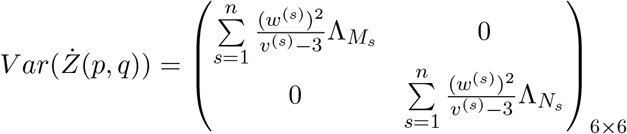

Substituting the variance covariance matrix of partial derivative of the random field by the FWHM using its relationship with |Λ|, we could get:

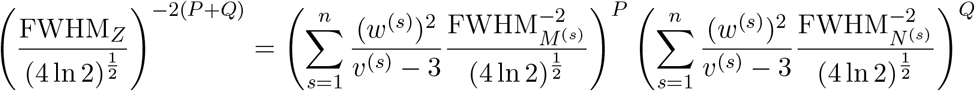

thus

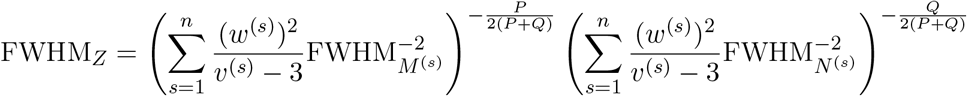

### F Supplement figures

**Figure 1:**
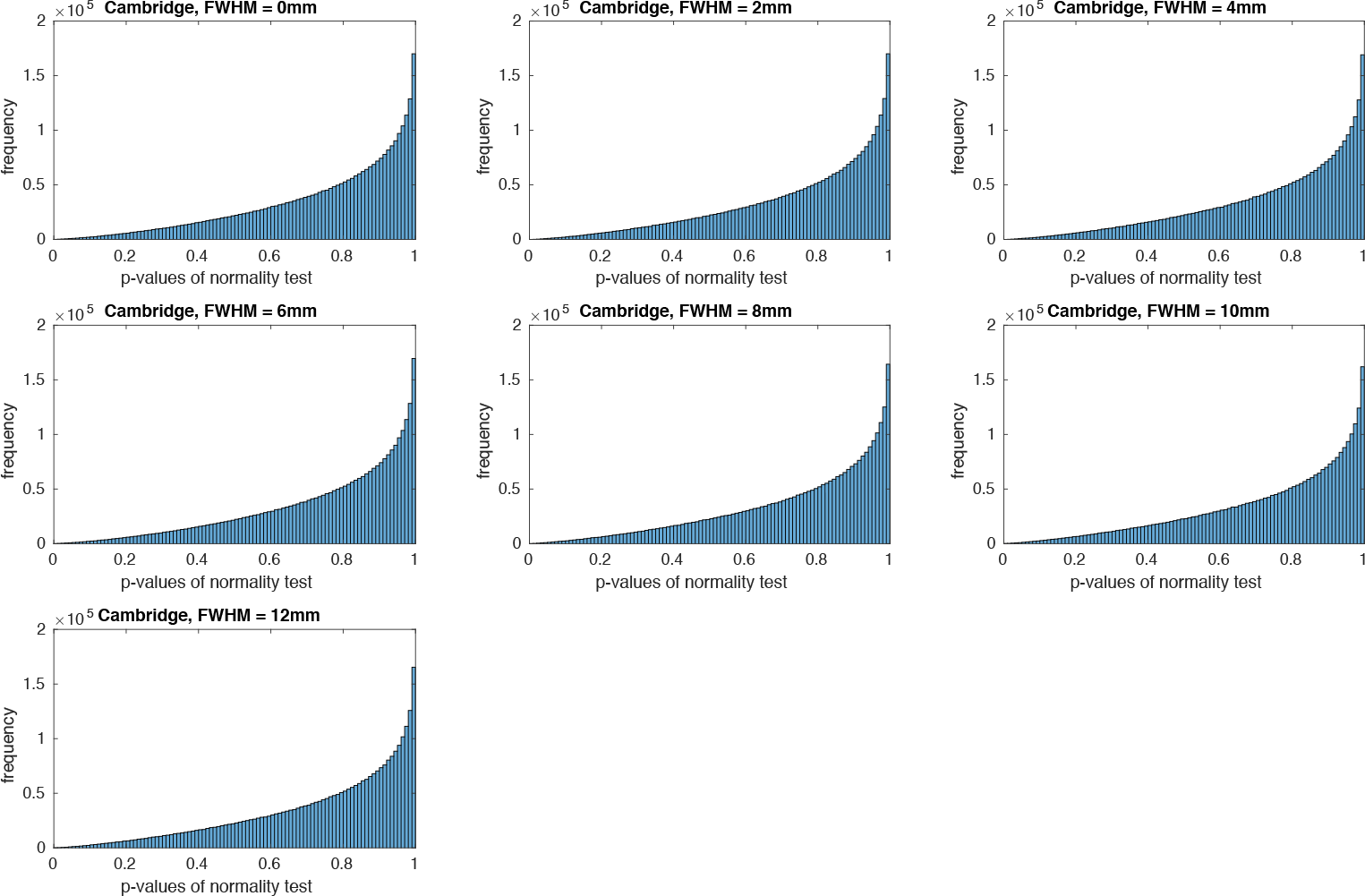
The p-values of normality tests in the Cambridge dataset.

**Figure 2:**
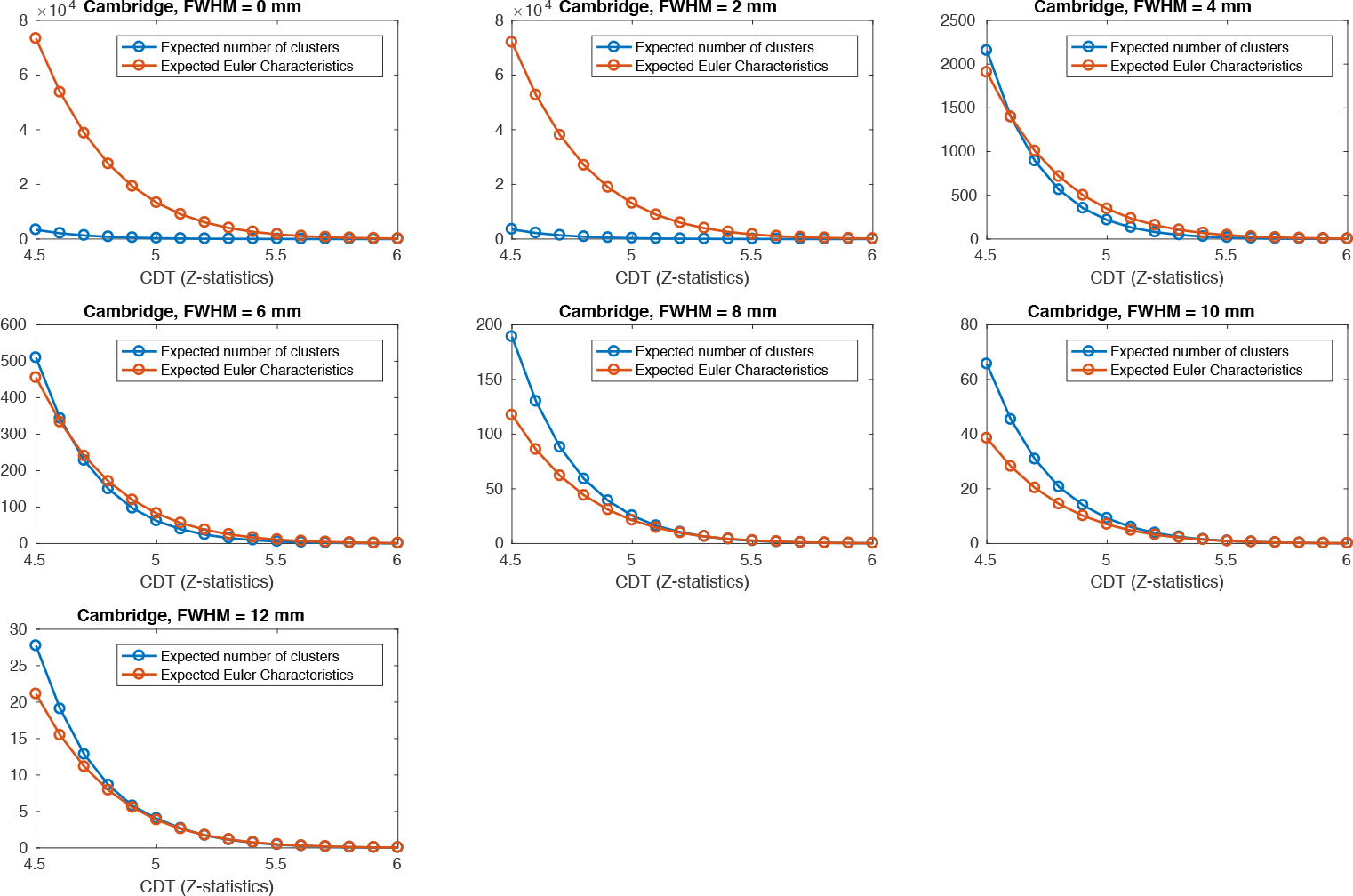
Comparing the expected Euler characteristics calculated by the Gaussian random field theory ith the observed expected number of clusters across different levels of CDT under different smoothness the Cambridge dataset.

**Figure 3:**
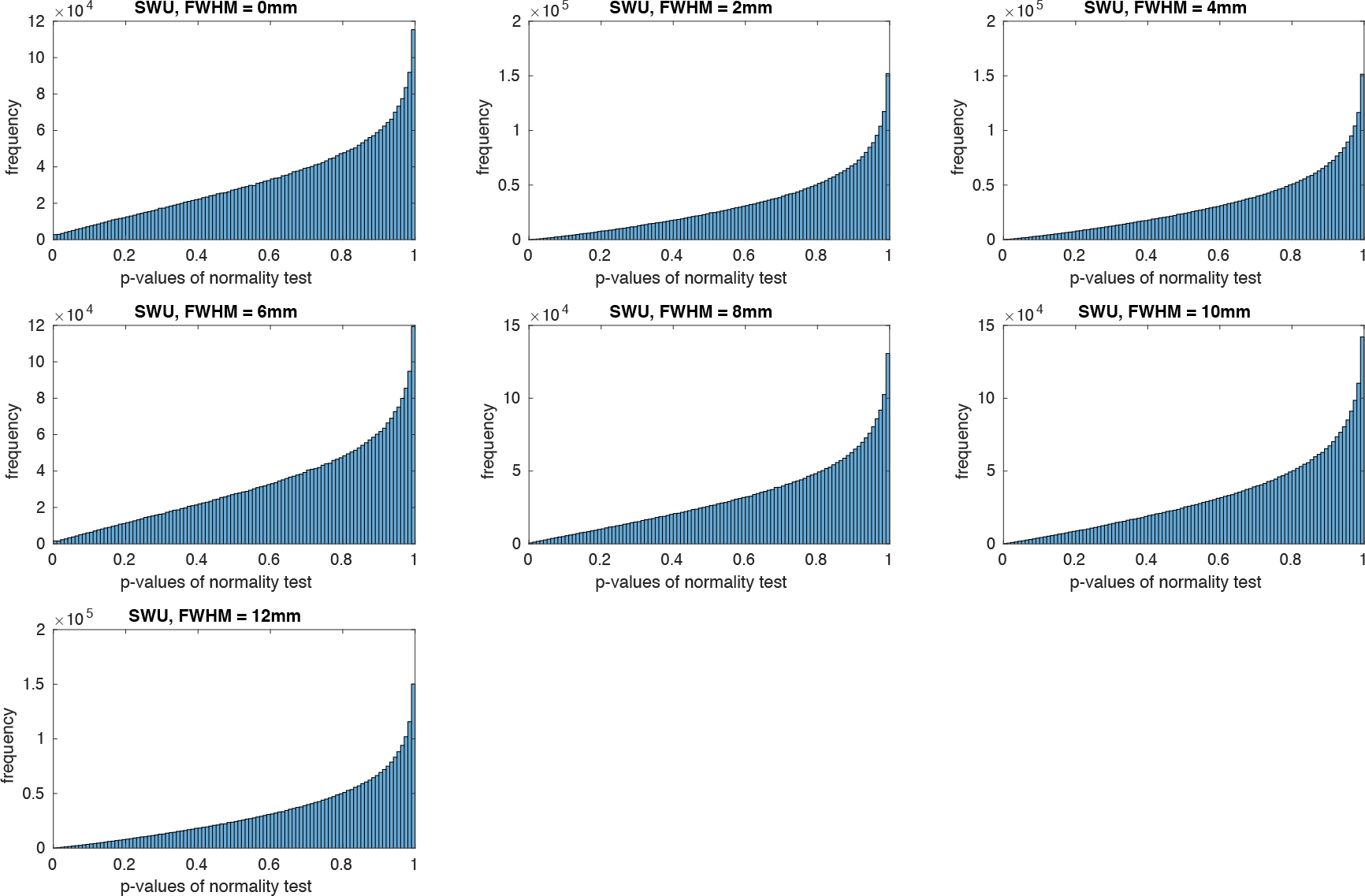
The p-values of normality tests in the Southwest University dataset.

**Figure 4:**
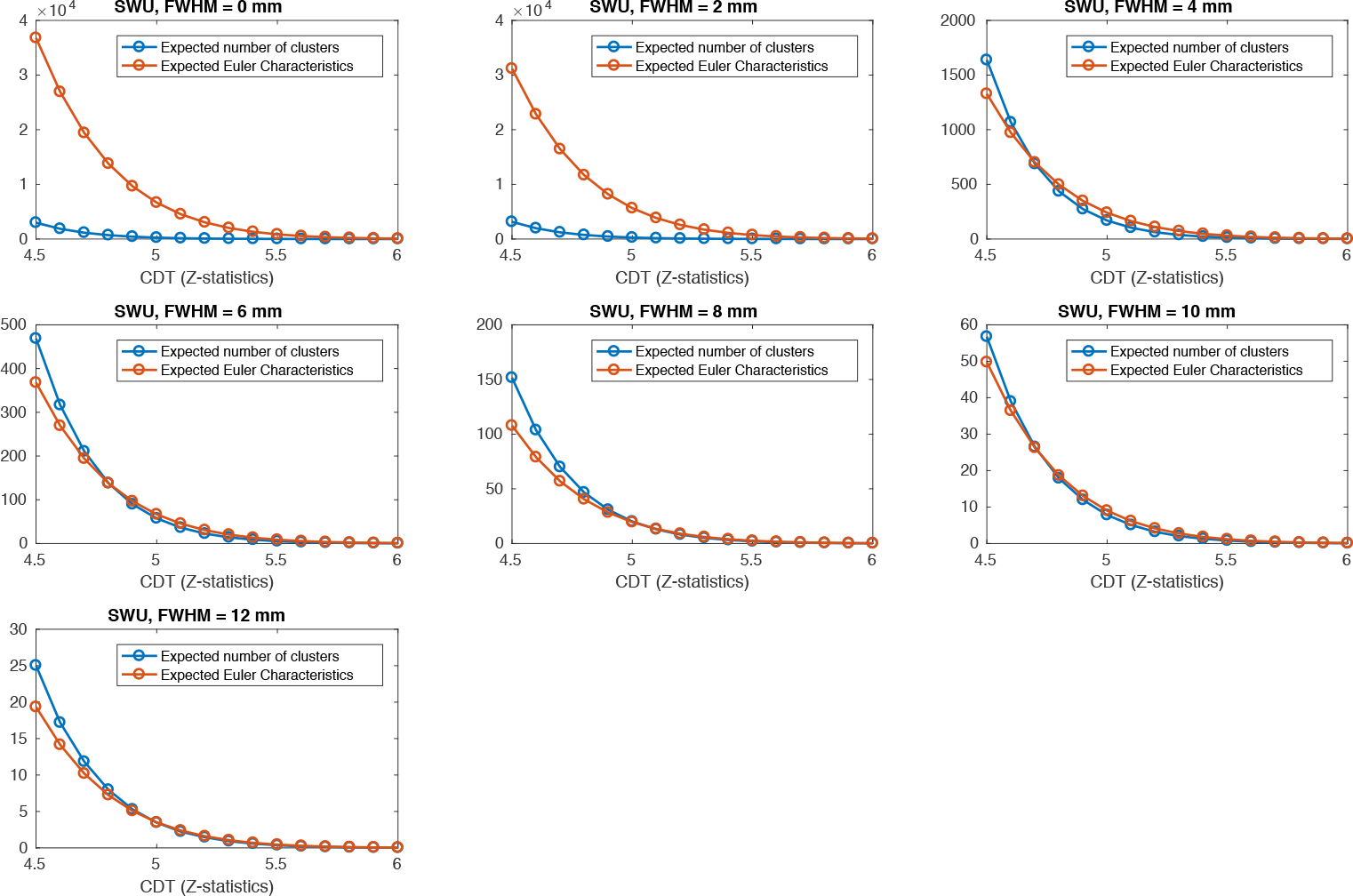
Comparing the expected Euler characteristics calculated by the Gaussian random field theory ith the observed expected number of clusters across different levels of CDT under different smoothness Southwest university dataset.

**Figure 5:**
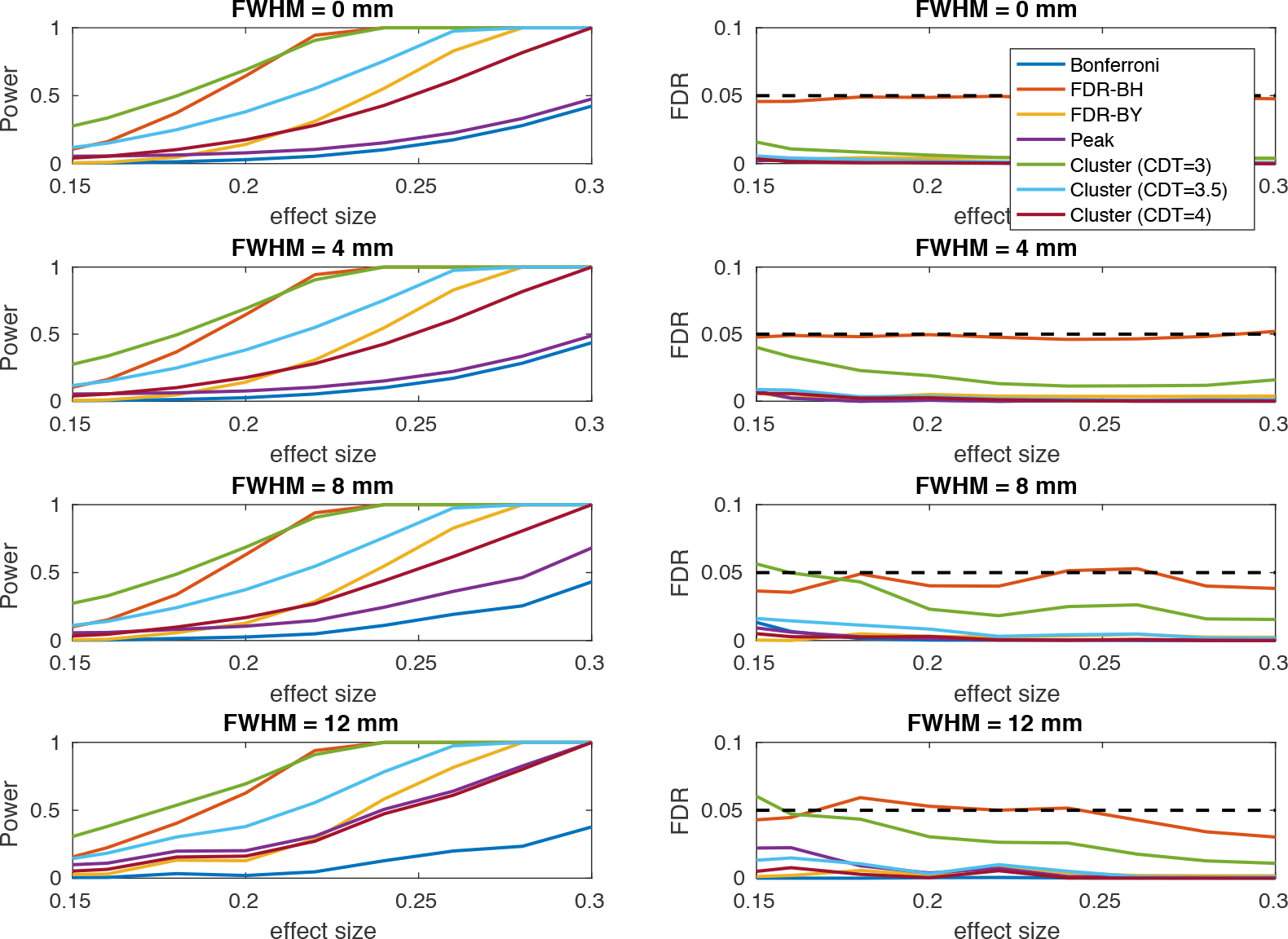
Similar to the Figure 8 in the main text, but this figure is generated by using different set of voxels.

**Figure 6:**
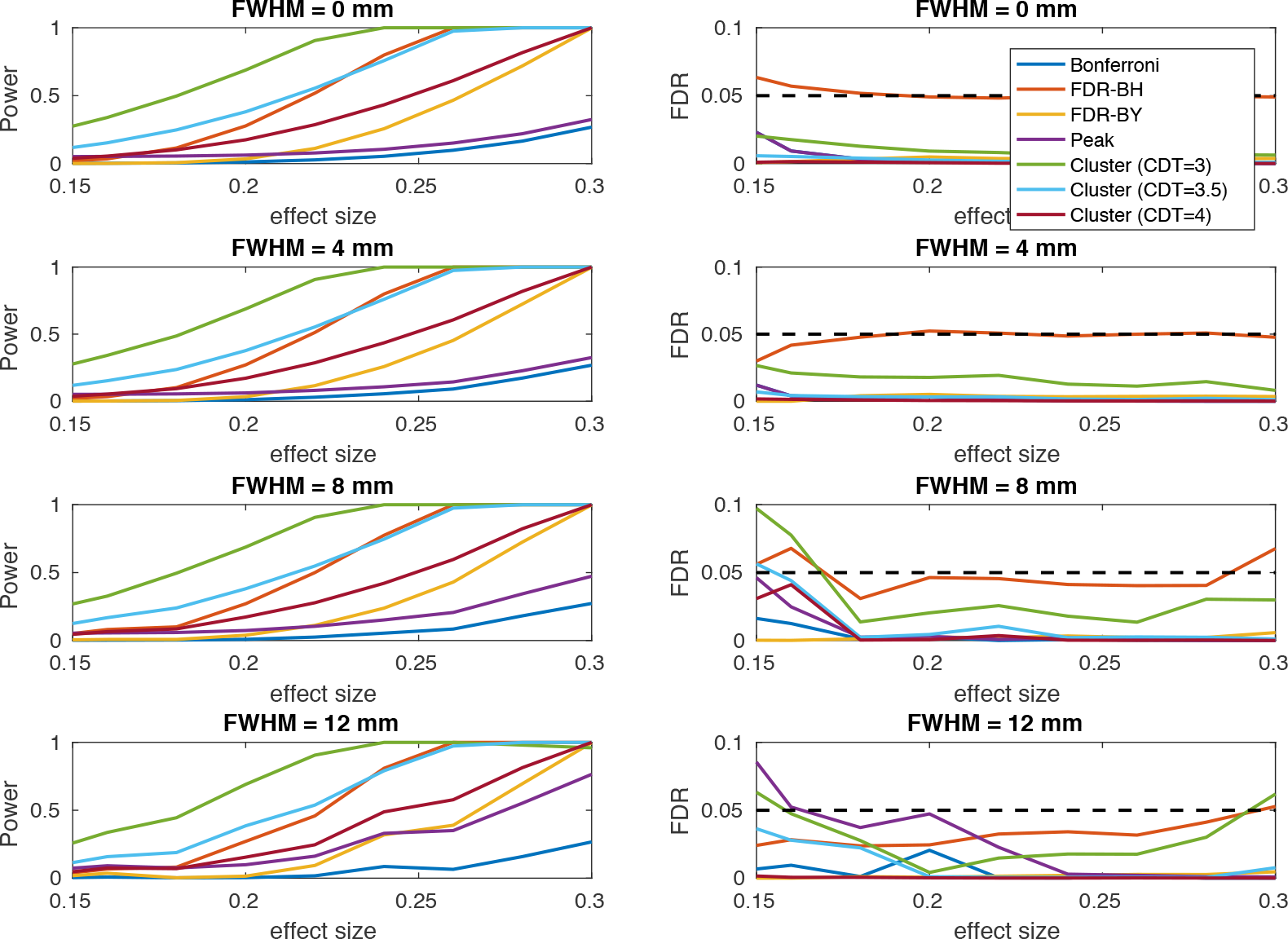
Similar to the Figure 8 in the main text, but this figure is generated by using different set of voxels.

**Figure 7:**
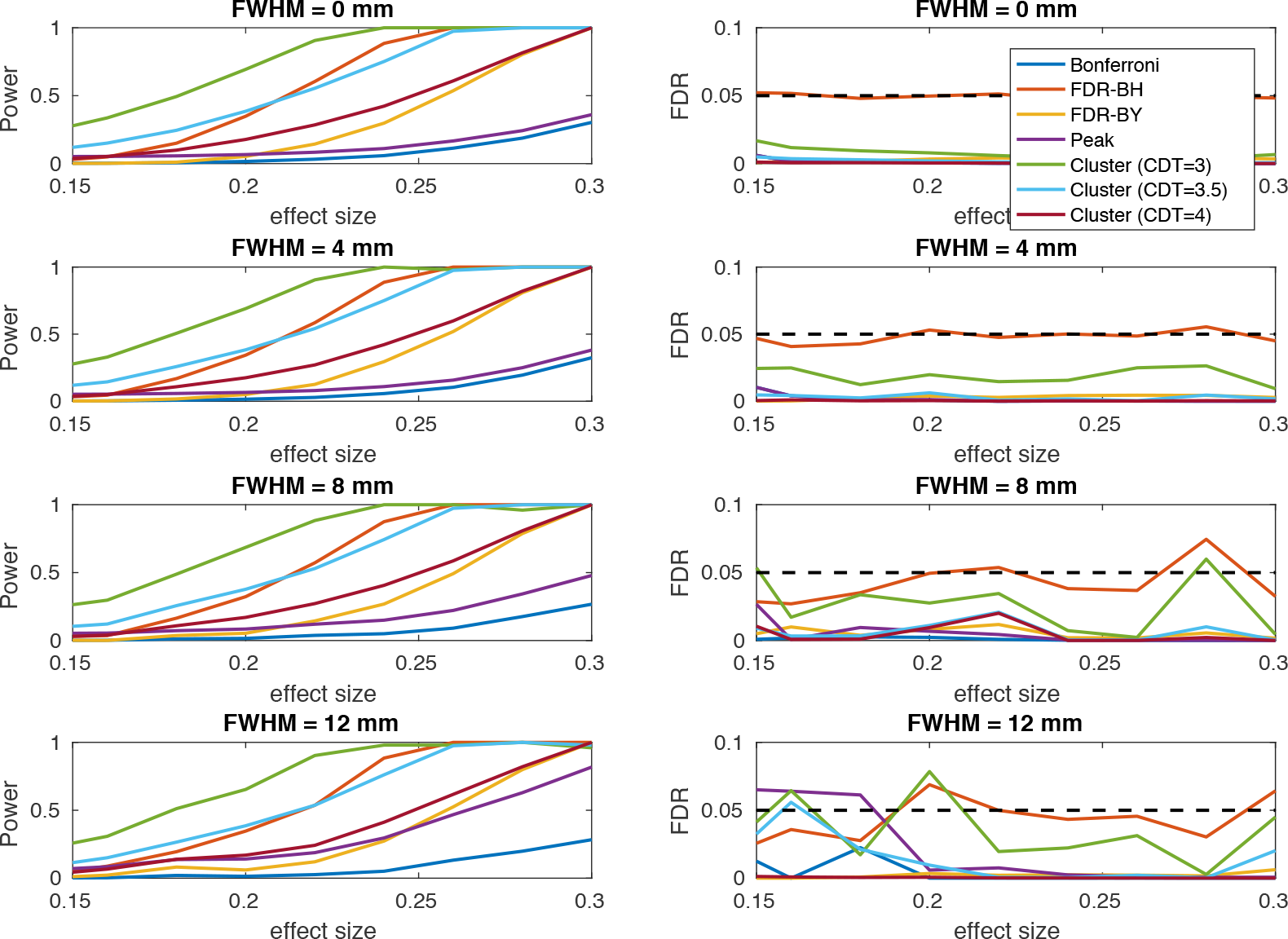
Similar to the Figure 8 in the main text, but this figure is generated by using different set of voxels.

### G Supplement tables

**Table 1:**
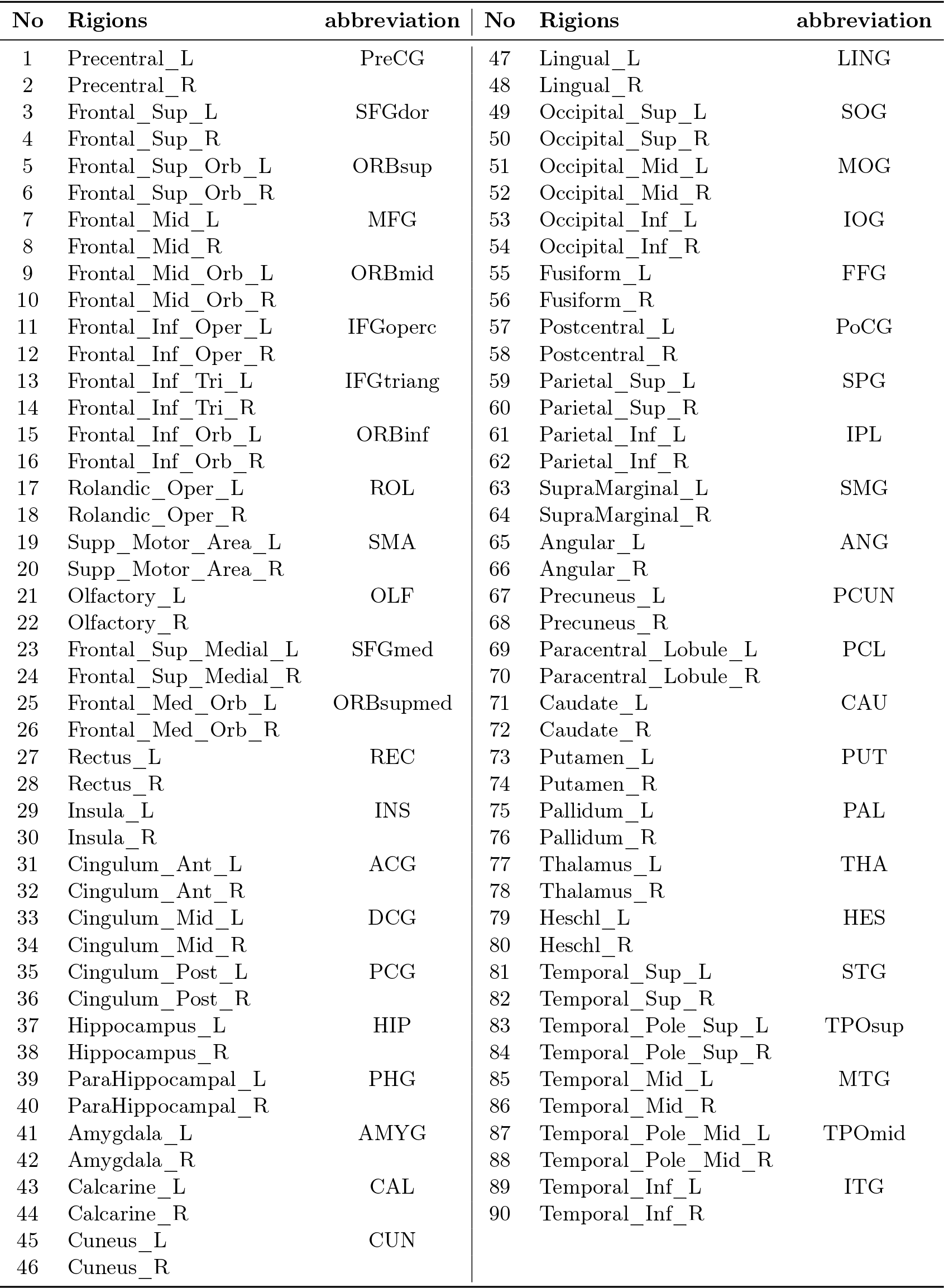
The names and abbreviations of anatomical regions of interest.

| No | Rigions | abbreviation | No | Rigions | abbreviation |
| --- | --- | --- | --- | --- | --- |
| 1 | Precentral_L | PreCG | 47 | Lingual_L | LING |
| 2 | Precentral_R |  | 48 | Lingual_R |  |
| 3 | Frontal_Sup_L | SFGdor | 49 | Occipital_Sup_L | SOG |
| 4 | Frontal_Sup_R |  | 50 | Occipital_Sup_R |  |
| 5 | Frontal_Sup_Orb_L | ORBsup | 51 | Occipital_Mid_L | MOG |
| 6 | Frontal_Sup_Orb_R |  | 52 | Occipital_Mid_R |  |
| 7 | Frontal_Mid_L | MFG | 53 | Occipital_Inf_L | IOG |
| 8 | Frontal_Mid_R |  | 54 | Occipital_Inf_R |  |
| 9 | Frontal_Mid_Orb_L | ORBmid | 55 | Fusiform_L | FFG |
| 10 | Frontal_Mid_Orb_R |  | 56 | Fusiform_R |  |
| 11 | Frontal_Inf_Oper_L | IFGoperc | 57 | Postcentral_L | PoCG |
| 12 | Frontal_Inf_Oper_R |  | 58 | Postcentral_R |  |
| 13 | Frontal_Inf_Tri_L | IFGtriang | 59 | Parietal_Sup_L | SPG |
| 14 | Frontal_Inf_Tri_R |  | 60 | Parietal_Sup_R |  |
| 15 | Frontal_Inf_Orb_L | ORBinf | 61 | Parietal_Inf_L | IPL |
| 16 | Frontal_Inf_Orb_R |  | 62 | Parietal_Inf_R |  |
| 17 | Rolandic_Oper_L | ROL | 63 | SupraMarginal_L | SMG |
| 18 | Rolandic_Oper_R |  | 64 | SupraMarginal_R |  |
| 19 | Supp_Motor_Area_L | SMA | 65 | Angular_L | ANG |
| 20 | Supp_Motor_Area_R |  | 66 | Angular_R |  |
| 21 | Olfactory_L | OLF | 67 | Precuneus_L | PCUN |
| 22 | Olfactory_R |  | 68 | Precuneus_R |  |
| 23 | Frontal_Sup_Medial_L | SFGmed | 69 | Paracentral_Lobule_L | PCL |
| 24 | Frontal_Sup_Medial_R |  | 70 | Paracentral_Lobule_R |  |
| 25 | Frontal_Med_Orb_L | ORBsupmed | 71 | Caudate_L | CAU |
| 26 | Frontal_Med_Orb_R |  | 72 | Caudate_R |  |
| 27 | Rectus_L | REC | 73 | Putamen_L | PUT |
| 28 | Rectus_R |  | 74 | Putamen_R |  |
| 29 | Insula_L | INS | 75 | Pallidum_L | PAL |
| 30 | Insula_R |  | 76 | Pallidum_R |  |
| 31 | Cingulum_Ant_L | ACG | 77 | Thalamus_L | THA |
| 32 | Cingulum_Ant_R |  | 78 | Thalamus_R |  |
| 33 | Cingulum_Mid_L | DCG | 79 | Heschl_L | HES |
| 34 | Cingulum_Mid_R |  | 80 | Heschl_R |  |
| 35 | Cingulum_Post_L | PCG | 81 | Temporal_Sup_L | STG |
| 36 | Cingulum_Post_R |  | 82 | Temporal_Sup_R |  |
| 37 | Hippocampus_L | HIP | 83 | Temporal_Pole_Sup_L | TPOsup |
| 38 | Hippocampus_R |  | 84 | Temporal_Pole_Sup_R |  |
| 39 | ParaHippocampal_L | PHG | 85 | Temporal_Mid_L | MTG |
| 40 | ParaHippocampal_R |  | 86 | Temporal_Mid_R |  |
| 41 | Amygdala_L | AMYG | 87 | Temporal_Pole_Mid_L | TPOmid |
| 42 | Amygdala_R |  | 88 | Temporal_Pole_Mid_R |  |
| 43 | Calcarine_L | CAL | 89 | Temporal_Inf_L | ITG |
| 44 | Calcarine_R |  | 90 | Temporal_Inf_R |  |
| 45 | Cuneus_L | CUN |  |  |  |
| 46 | Cuneus_R |  |  |  |  |

